# PD-1 blockade drug holiday improves exhausted progenitor CD8 T cell (Tpex) reinvigoration by avoiding Tpex adaptive resistance

**DOI:** 10.64898/2026.05.14.725199

**Authors:** Shin Foong Ngiow, Ryan P. O’Connell, Sasikanth Manne, Yinghui Jane Huang, Josephine R. Giles, Victor Alcalde, Divij Mathew, Max Klapholz, Kito Nzingha, Zeyu Chen, Amy E. Baxter, Kunal P. Patel, Jennifer E. Wu, Ryan P. Staupe, Mohammed-Alkhatim A. Ali, Allison R. Greenplate, David Weiner, E. John Wherry

## Abstract

Blocking the programmed cell death 1 (PD-1) pathway using monoclonal antibodies reinvigorates exhausted T cells (Tex), enhancing control of chronic viral infections and cancer. Considerable effort has focused on evaluating different PD-1 blockade agents in preclinical and clinical cancer settings, but relatively little information exists on how to optimize the pharmacodynamic effects of PD-1 pathway blockade on reinvigorating Tex. To address this question, we performed longitudinal tracking of Tex reinvigoration during chronic infection with lymphocytic choriomeningitis virus (LCMV) following different regimens of PD-1 blockade. We compared single-cycle (2 weeks of treatment), long-term continuous PD-1 pathway blockade (i.e. 3 months), or blockade followed by a drug holiday and then re-blockade (intermittent treatment). These studies revealed little benefit of continuous versus single-cycle PD-1 blockade, with both resulting in a single peak of Tex reinvigoration and similar effects on viral replication. In contrast, intermittent blockade resulted in a new cycle of secondary Tex reinvigoration upon redosing after a washout and this secondary Tex reinvigoration improved disease control. Mechanistically, long-term blockade eroded the ability of Tex progenitor cells (Tpex) to give rise to downstream, more functional Tex intermediate (Tex-Int) progeny, whereas the drug holiday restored this Tpex proliferative and differentiation capacity. Tpex from long-term treated mice showed evidence of adaptive resistance and additional layers of negative regulation, including sustained expression of the inhibitory receptor CD22. Indeed, co-blockade of PD-1 and CD22 using combination antibodies or bispecific antibody approaches improved disease control and reinvigoration of Tex. These data have implications for clinical immune pharmacodynamics of PD-1 blockade and provide insights into the biology of Tex reinvigoration.

**One Sentence Summary:** Modifying the immunopharmacology of PD-1 blockade reveals a benefit of a drug holiday and identifies mechanisms of Tex progenitor deficiency provoked by prolonged loss of PD-1 signals including the inhibitory receptor CD22.

## Main

T cell exhaustion is a state of T cell differentiation associated with impaired effector functions and poor control of chronic infections and cancer ^1^. Sustained expression of inhibitory receptors (IRs) including PD-1 by Tex limits their ability to control persisting infections and tumors ^2^. However, Tex can be partially reinvigorated by PD-1 pathway blockade boosting proliferation, cytokine production and cytotoxicity, improving disease control ^2^. This Tex reinvigoration by PD-1 pathway blockade works by targeting Tex progenitor cells (Tpex) causing these cells to proliferate and differentiate into downstream, more differentiated and cytotoxic effector-like Tex intermediate (Tex-Int) and terminally exhausted (Tex-Term) cells ^3–10^. Indeed, Tex reinvigoration is central to the clinical response to PD-1 blockade in human cancers ^11–15^. PD-1 pathway blockade drugs are approved in 20+ different human cancer settings and with thousands of active clinical trials testing PD-1/L1 blockade agents in immune oncology settings ^16–18^.

Despite the clinical success of PD-1 pathway blockade key questions remain about the optimal strategies for using antibodies targeting PD-1 (or PD-L1) to provoke reinvigoration of Tex. For example, data from mice ^19^ and humans ^11,12^ indicate that PD-1 pathway blockade provokes only a single, transient, peak of Tex reinvigoration despite, at least in humans, the continuous blockade of PD-1. These observations suggest that although the pharmacodynamics of the anti-PD-1 (or anti-PD-L1) biologic drug used clinically are well-defined, the immune pharmacodynamics of the reinvigorated Tex contributing to disease control are at least partially disconnected from treatment regimen. This concept of anti-PD-1 as the “prodrug” and reinvigorated Tex as (part of) the *in vivo* drug (i.e. modified immune system), provoke questions about how anti-PD-1 treatment regimens could be optimized for more effective Tex reinvigoration. Several challenges exist to addressing these questions including the use of species-mismatched PD-1 blocking antibodies in most pre-clinical studies that result in anti-drug antibodies and the paucity of preclinical models that can be studied for prolonged periods of time to assess durability of Tex reinvigoration. These issues further limit the study of adaptive resistance mechanisms that may be induced intrinsically in Tex due to continuous PD-1 blockade. Thus, it has been unclear whether prolonged PD-1 blockade is the best approach to achieve optimal Tex reinvigoration.

In this study we addressed these questions by using chronic LCMV infection where T cell exhaustion is achieved at a steady state for months to years ^7,19–23^. We then also employed a mouse anti-mouse PD-1 blockade antibody to test and compare short-term, long-term, and intermittent (Drug Holiday) PD-1 blockade and ask whether these different treatments resulted in differences in the magnitude and/or durability of reinvigorated antigen-specific Tex. Nearly all preclinical data for PD-1 pathway blockade has been generated using a treatment regimen where blocking antibodies are administered every 1-3 days, but only for a short 1-2 week “single-cycle” of blockade. In contrast, human PD-1 blockade is typically continuous or long-term (i.e. often many months to >1 year). We therefore first examined these two scenarios in the LCMV clone 13 infection model where long-term anti-PD-1 blockade could be achieved because of the establishment of a long-term disease steady-state. Thus, we compared antigen-specific CD8 Tex reinvigoration during different PD-1 blockade regimens of Short-Term (ST; two weeks of PD-1 blockade starting at day 22 (d22) of chronic infection; i.e. dosing every 3 days for 2 weeks) versus Long-Term (LT; continuous PD-1 blockade for 12 weeks starting on d22, dosing every 3 days) (**Fig. 1a**). Following either blockade regimen, Tex were initially reinvigorated and the frequency and number of LCMV specific TCR transgenic P14 cells increased robustly in the peripheral blood of anti-PD-1 treated mice compared to Control (Ctrl) mice, peaking ∼2 weeks following initiation of treatment. However, for both regimens this numerical reinvigoration returned to baseline by 6-8 weeks post treatment initiation. In the case of the LT treatment this return to baseline occurred despite the continued delivery of anti-PD-1 antibody. For the LT group there was perhaps an advantage in percentage of responding cells at the d65 time point, but not in number of reinvigorated Tex/10^6^ PBMC (**Fig. 1b**). Indeed, the reinvigorated Tex response in both the ST and LT mice subsequently contracted with similar kinetics (**Fig. 1b**). Moreover, 12 weeks after the start of PD-1 blockade therapy, the frequency and number of P14 cells in spleen and liver, were comparable between ST and LT mice, (**Fig. 1c,d**). Similar results were obtained analyzing the endogenous, non-TCR transgenic D^b^GP_33_-tetramer+ CD8 T cell response (**Extended Data Fig. 1a**). Ki67 expression was also similar between these groups at the week 12 time point (**Fig. 1e; Extended Data Fig. 1b**). These data suggest that prolonged PD-1 blockade provided no additional benefit to further expand nor sustain the proliferative or numerical response of reinvigorated Tex compared to short-term PD-1 blockade.

**Fig. 1.**
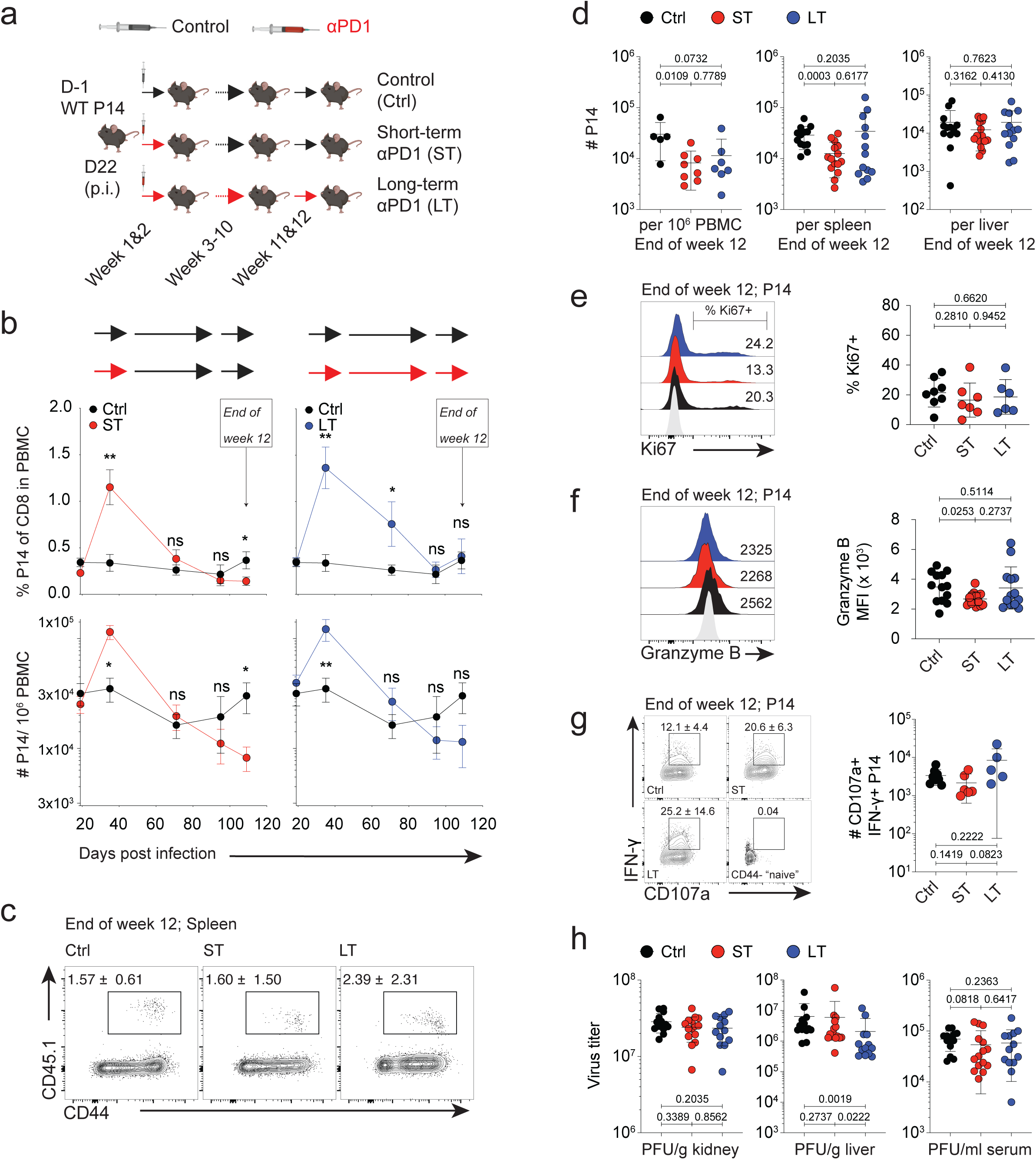
Comparison of reinvigorated Tex responses between transient and long-term PD-1 blockade. Groups of LCMV clone 13-infected B6 mice containing adoptively transferred P14 cells were treated with the indicated PD-1 blockade regimens. (a) Experimental design. (b) Longitudinal analysis of the frequency (upper panel) and number (lower panel) of P14 cells in blood of LCMV clone 13-infected mice with the indicated PD-1 blockade regimen. Data shown are presented as mean ± SEM. Black and red arrows shown above individual graphs represent control (Ctrl) or PD-1 blockade, respectively, as illustrated in (a). (c) Representative plots for spleen P14 cells are shown, with frequency depicted as mean ± SD. (d) Numbers of P14 cells in blood, spleen, and liver. (e) Representative overlaid histograms and frequencies for Ki67+ cells as a fraction of P14 cells are shown. (f) Representative overlaid histograms and MFIs for Granzyme B of P14 cells are shown. (g) Flow cytometry plots (concatenated P14 cells), frequencies (depicted in mean ± SD) and numbers of CD107a+IFN-γ+ P14 cells in spleen are shown. (h) LCMV viral load in kidney, liver, and serum of mice following different PD-1 blockade regimens are shown. Grey histogram in (e) and (f): gated on CD44– naïve cells. Each dot in (d-h) represents an individual mouse, and error bars represent SD. Statistical significance between groups was determined by two-tailed Mann-Whitney test (* P < 0.05; ** P < 0.01; ns, not significant). Data shown in (b), (d) PBMC P14 (n = 5-8 per group), (e) (n = 6-8 per group), and (g) (n = 5-8 per group) are representative of two independent experiments. Data shown in (d) spleen and liver P14, (f) and (h) (n =13-15 per group) are pooled from two independent experiments.

In addition to a proliferative response, PD-1 blockade also reinvigorates effector activity, but previous studies showed that by 5 months after cessation of a ST treatment regimen, functional improvements such as increased effector molecule expression had returned to baseline ^19^. Indeed, Granzyme B expression was slightly decreased in ST versus Ctrl-treated P14 cells at 12 weeks (**Fig. 1f; Extended Data Fig. 1c**). However, even in the LT group receiving continuous PD-1 blockade, Granzyme B expression was unchanged compared to Tex from Ctrl-treated mice. Similarly, the number of P14 cells with the ability to degranulate (CD107a) and produce IFN-γ, following *ex vivo* peptide restimulation, was similar between Ctrl, ST and LT treatment groups at the week 12 time point (**Fig. 1g**). These data suggest that continuous blockade of the PD-1 pathway did not result in sustained improvement in effector functions and was no better than short-term blockade at sustaining reinvigoration of Tex.

A major benefit of PD-1 pathway blockade is improved disease control, including reduced viral burden during LCMV clone 13 infection ^24^. Immediately after a two-week treatment regimen viral load is reduced (^19,24^ ;data not shown). However, 10 weeks later there was no reduction in viral load in kidney, liver, or serum between ST and Ctrl-treated mice (**Fig. 1h**), indicating a viral rebound after initial anti-PD-1-induced viral control. After 12 weeks of treatment the viral burden in liver of LT mice was slightly reduced compared to Ctrl mice (**Fig. 1h *middle panel***), consistent with a higher number of endogenous D^b^GP_33_-tetramer+ Tex in the liver of LT mice (**Extended Data Fig. 1a**). However, despite 12 weeks of continued PD-1 pathway blockade there was also no reduction in viral load in the kidney or serum between LT and Ctrl-treated mice (**Fig. 1h**), suggesting that the initial systemic benefit of viral control was lost over time. Overall, the LT treatment group displayed relatively little benefit in viral control across all tissues compared to either the ST or Ctrl-treated group (**Fig. 1h *left* and *right panels***). In agreement with the overall disease control effect observed in chronic LCMV, there was no difference in disease control in MCA1956 tumor-bearing mice after short-term (d8-d20 dosing every 4 days) versus long-term (d4-d84 dosing every 4 days) PD-1 blockade regimens (**Extended Data Fig. 2**). Together, these data indicate that Tex reinvigoration and disease burden are only minimally improved by long-term PD-1 blockade compared to transient blockade and are consistent with data from humans indicating that most of the immunological response to PD-1 pathway blockade occurs shortly after treatment initiation ^11–15^.

We next tested whether a different PD-1 blockade regimen would improve the therapeutic effect of reinvigorating Tex. We asked whether cycles of PD-1 blockade followed by rest from anti-PD-1 (i.e. a drug washout phase) would have any benefit. Such a “drug holiday” treatment approach has been difficult to achieve in preclinical models because tumor models are typically progressive, preventing long-term experiments needed for drug washout (**Extended Data Fig. 2**) and most preclinical models have used rat antibodies to the PD-1 pathway resulting in the induction of species-specific anti-drug antibodies. Similarly, in humans the prolonged cell surface bound half-life of many months for some clinical PD-1 pathway drugs ^25^ makes such a drug washout and retreatment approach challenging. Thus, we used the chronic LCMV infection model with steady state viremia and T cell exhaustion, and a mouse anti-mouse PD-1 blocking antibody to address this question. We tested whether a therapy washout period (Drug Holiday) would permit a second Tex reinvigoration event by anti-PD-1 retreatment. We compared the Tex response in mice that received the following PD-1 blockade regimens: early ST treatment (ST^early^; same experiment as described in **Fig. 1a**), Drug Holiday (two separate periods of two weeks of PD-1 blockade at weeks 1-2 and 11-12 dosing every 3 days, separated by an eight-week washout period during weeks 3-10), late ST treatment (ST^late^; two weeks of PD-1 blockade in weeks 11-12 dosing every 3 days) (**Fig. 2a**). Unlike mice that received continuous blockade in **Fig. 1**, mice receiving the Drug Holiday regimen experienced a second burst of Tex reinvigoration following re-blockade from weeks 11-12 (**Fig. 2b**) suggesting a benefit of a drug washout phase. The magnitude of P14 reinvigoration at this late time point was similar between the Drug Holiday and the ST^late^ treatment (**Fig. 2b**). Compared to ST^early^ mice (**Fig. 2c,d**) or the LT mice (**Fig. 2e**), Drug Holiday mice had an increased number of P14 cells in the peripheral blood and liver. This numerical reinvigoration of Tex was nearly as robust for the Drug Holiday group receiving a second round of PD-1 blockade as for the ST^late^ mice receiving PD-1 blockade for the first time (**Fig. 2c,d**), though Ki67 was higher in the ST^late^ group (**Fig. 2f**). A Drug Holiday also improved responses in non-TCR transgenic CD8 T cells, with increased Ki67 and a trend towards a numerical increase, consistent with a second round of reinvigoration (**Extended Data Fig. 3a,b**). Granzyme B expression was increased in the reinvigorated Tex in both the Drug Holiday and ST^late^ treatment regimens, though the magnitude of increase was slightly higher for the ST^late^ group (**Fig. 2g; Extended Data Fig. 3c**). Similarly, the Drug Holiday regimen improved other effector functions including the number of P14 capable of degranulation and IFN-γ production (**Fig. 2h**). Collectively, these data indicate that an intermittent PD-1 blockade regimen allows a second round of anti-PD-1 mediated Tex reinvigoration and re-expansion.

**Fig. 2.**
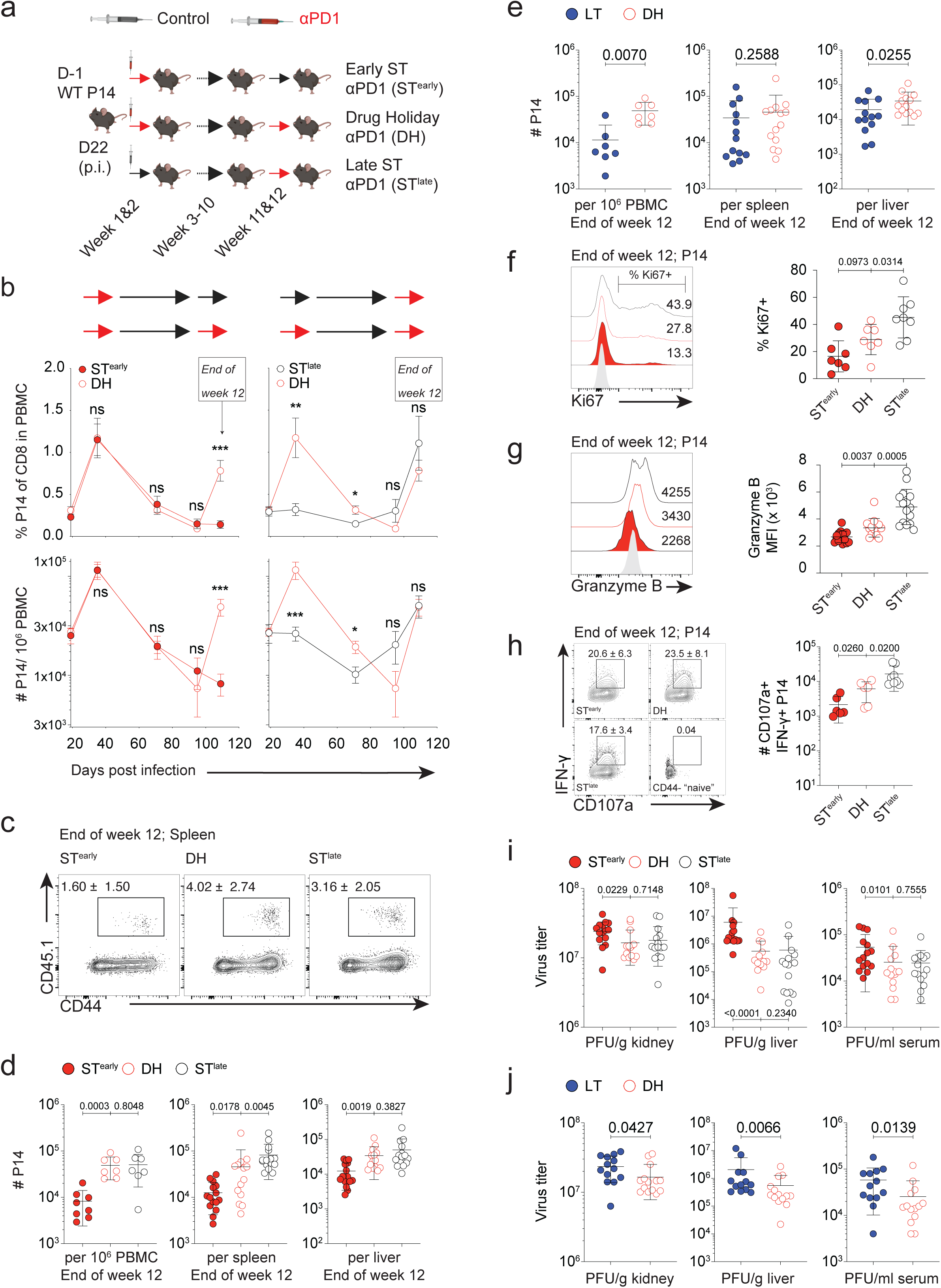
Intermittent PD-1 blockade permits secondary Tex reinvigoration. Groups of LCMV clone 13-infected B6 mice containing adoptively transferred P14 cells were treated with the indicated PD-1 blockade regimen. (a) Experimental design. Note: Treatment groups shown in Fig. 2a were performed concurrently with those in Fig. 1a. (b) Longitudinal analysis of the frequency (upper panel) and number (lower panel) of P14 cells in blood of LCMV clone 13-infected mice with the indicated PD-1 blockade regimen. Data shown are presented as mean ± SEM. Black and red arrows shown above individual graphs represent control or PD-1 blockade, respectively, as illustrated in (a). (c) Representative plots for spleen P14 cells are shown, with frequency depicted as mean ± SD. (d and e) Numbers of P14 cells in blood, spleen and liver. (f) Representative overlaid histograms and frequencies for Ki67+ cells as a fraction of P14 cells are shown. (g) Representative overlaid histograms and MFIs for Granzyme B of P14 cells are shown. (h) Flow cytometry plots (concatenated P14 cells), frequencies (depicted in mean ± SD) and numbers of CD107a+IFN-γ+ P14 cells in spleen are shown. (i and j) LCMV viral load in kidney, liver, and serum of mice following different PD-1 blockade regimens are shown. Grey histogram in (f) and (g): gated on CD44– naïve cells. Each dot in (d-j) represents an individual mouse, and error bars represent SD. Statistical significance between groups was determined by two-tailed Mann-Whitney test (* P < 0.05; *** P < 0.001; ns, not significant). Data shown in (b), (d and e) PBMC P14, (f), and (h) (n = 6-8 per group) are representative of two independent experiments. Data shown in (d and e) spleen and liver P14, (g) and (i and j) (n = 13-15 per group) are pooled from two independent experiments.

The effects of Tex reinvigoration in the Drug Holiday group suggested potential benefits on disease control. Indeed, in contrast to the ST^early^ or LT treatment groups, the secondary Tex reinvigoration in the Drug Holiday treatment group resulted in improved control of viral replication in the serum, liver and kidney compared to the ST^early^ group (**Fig. 2i**) or the LT group (**Fig. 2j**). This viral control in the Drug Holiday group was similar to that achieved in the ST^late^ treatment group receiving anti-PD-1 therapy for the first time (**Fig. 2i**). These observations reveal the potential benefit of an intermittent PD-1 blockade regimen with improved Tex reinvigoration and disease control.

To begin to explore the underlying mechanisms of LT and Drug Holiday treatment regimens, we asked whether single round, continuous or intermittent PD-1 blockade regimens had distinct impacts on the differentiation state of responding Tex using high-dimensional cytometry at the end of all treatments (i.e. d110, end of week 12). Expression patterns of IRs or LY108/TCF-1 have been used to define subsets of Tex with different biological properties ^3,5–7,20,26^. Tex-Term have higher IR expression, low LY108/TCF-1 and fail to respond to anti-PD-1 whereas TCF-1-expressing Tpex respond to PD-1 pathway blockade ^5–7^. This pattern of Tex differentiation can be further resolved using CX3CR1 to identify LY108–CX3CR1+ Tex-Int with improved effector activity, in addition to a LY108+CX3CR1– Tpex and a LY108–CX3CR1– Tex-Term population ^9,10^. LY108 and CD69 expression also identifies two distinct LY108+ (CD69+ and CD69–) Tpex populations, a LY108–CD69– Tex-Int population, and a LY108–CD69+ Tex-Term population ^8^. Thus, we investigated whether different anti-PD-1 treatment approaches altered Tex subset biology by assessing the distribution of Tex subsets based on LY108, CX3CR1 and/or CD69. We first focused on the “effector-like” Tex-Int subsets that may be important for therapeutic efficacy and disease control following PD-1 pathway blockade ^8–10^. Compared to Tex from Ctrl-treated mice, on d110 there were fewer Tex-Int in the ST^early^ and LT treatment groups (**Fig. 3a,b; Extended Data Fig. 4a,b, 5a,b,** and **6a,c,d, f**). In contrast, Tex-Int were expanded following PD-1 blockade in the ST^late^ and Drug Holiday treatment groups (**Fig. 3a,b; Extended Data Fig. 4a,b, 5a,b,** and **6a,c,d,f**), consistent with increased Granzyme B expression (**Fig. 2g**) and the other benefits of intermittent treatment (**Fig. 2i**). In addition to changes in the effector-like Tex-Int subset, the frequency of the LY108–CD69+ Tex-Term subset was increased in the Drug Holiday and LT groups, but not in the ST^late^ group (**Extended Data Fig. 4a****, 5b,** and **6d**). These proportional changes in LY108– Tex subsets also resulted in an increase in the Tex-Int to Tex-Term ratio in the Drug Holiday group compared to the LT group (**Fig. 3c; Extended Data Fig. 5c,** and **6b,e**). Despite a trend for a higher frequency of the Tex-Term subset in the ST^early^ and LT treatment groups compared to the Ctrl, the number of Tex-Term was not different between these groups (**Fig. 3b; Extended Data Fig. 4b,** and **6c,f**). Thus, whereas ST^early^ or LT treatment was associated with a decrease in the CX3CR1+ effector-like Tex-Int population, the Drug Holiday treatment strategy increased the frequency of this Tex subset, similar to that observed in the ST^late^ treatment group.

**Fig. 3.**
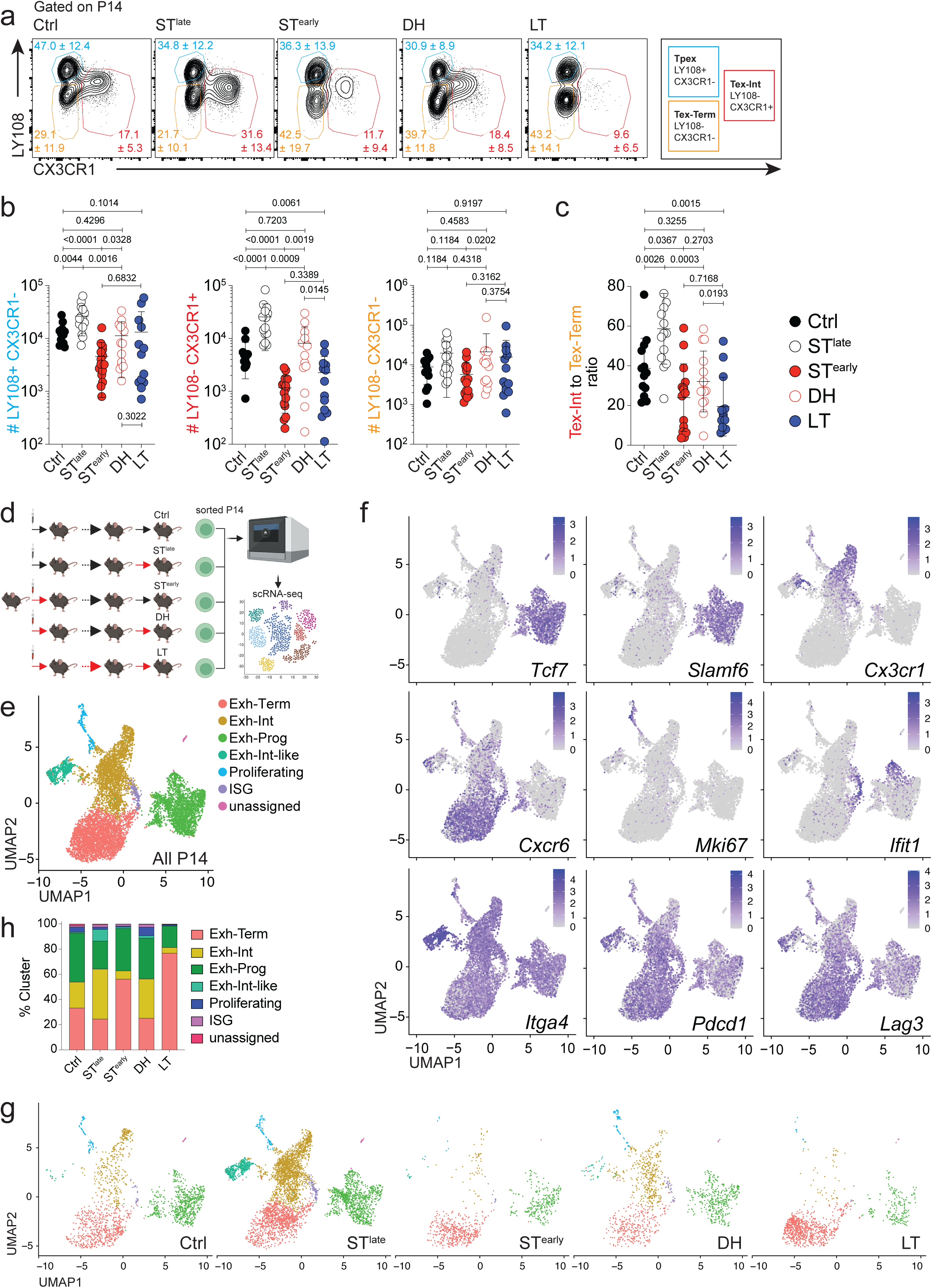
Comparison of P14 Tex subsets following different PD-1 blockade regimens. (a) Representative flow cytometry plots, frequencies (depicted in mean ± SD; See Extended Data Fig. 5a) and (b) numbers of LY108 CX3CR1 Tex subsets in the spleen following different PD-1 blockade regimens are shown. (c) P14 Tex-Int (LY108–CX3CR1+) to Tex-Term (LY108–CX3CR1–) ratios in the spleen following different PD-1 blockade regimens are shown. (d) Experimental schematics of scRNA-seq following different PD-1 blockade regimens as shown in Fig. 1a and 2a. (e) UMAP of all P14 Tex are colored by cluster. (f) Expression of the indicated genes in UMAP from (e). (g) UMAP of P14 Tex colored by cluster, following different PD-1 blockade regimens are shown. (h) Proportional changes in Tex cluster summarizing (g). (b and c) Each dot represents an individual mouse, and error bars represent SD. Statistical significance between groups was determined by two-tailed Mann-Whitney test. (a, b, and c) Data shown are pooled from two independent experiments (n = 13-15 per group). (e, f, and g) Dots represents individual P14 Tex cells.

We hypothesized that the different treatment strategies might impact the quantity of LY108+ Tpex. At d110, the numbers of LY108+ Tpex were lower in the ST^early^ compared to the Control treatment group (**Fig. 3b; Extended Data Fig. 4b**), consistent with a role for PD-1 in preserving Tpex populations ^27,28^. Although there was more variability in the LT group, prolonged anti-PD-1 treatment did not increase the number of LY108+ Tpex populations compared to the ST^early^ treatment group (**Fig. 3b; Extended Data Fig. 4b,** and **6c,f**). In contrast, the Drug Holiday regimen resulted in modestly improved numbers of Tpex compared to ST^early^ treatment (**Fig. 3b; Extended Data Fig. 4b**, **and 6c,f**). Thus, consistent with the idea that permanent loss of PD-1 signals can compromise Tpex biology over time ^27,28^, an intermittent treatment regimen appeared to have less detrimental effects than the LT treatment schedule.

These data are consistent with the notion that persistently stimulated CD8 T cells tune TCR signaling threshold, in part using PD-1, to establishing a host-pathogen (or host-tumor) stalemate ^29^. Indeed, PD-1 stabilizes the formation and stability of the TCF-1+ Tex precursor subset at early time points ^27^. Thus, we hypothesized that different anti-PD-1 blocking regimens might provoke distinct compensatory PD-1 expression patterns specifically by Tpex populations. Using a non-competitive staining strategy to evaluate PD-1 expression (**Extended Data Fig. 7**), the LY108+CX3CR1– Tpex from the ST^early^ treatment group expressed slightly more PD-1 than Tpex in the Ctrl group despite the cessation of anti-PD-1 antibodies for 10 weeks (**Extended Data Fig. 4c**). In ST^late^, Drug Holiday and LT mice, PD-1 expression was even higher on the LY108+CX3CR1– Tpex (**Extended Data Fig. 4c**). A similar pattern was observed even when assessing heterogeneity within the Tpex population ^5–10,26^ (**Extended Data Fig. 4d**). The increase in PD-1 expression in the ST^late^ and Drug Holiday treatment groups is consistent with reinvigoration. The elevated Tpex PD-1 expression in LT mice that do not experience sustained reinvigoration, however, suggests an induction of an adaptive compensatory mechanism to restrain Tex proliferation from continuous PD-1 blockade.

To interrogate the mechanisms that regulate Tex reinvigoration from different PD-1 blockade treatment regimens, we performed single-cell RNA-sequencing (scRNA-seq) of antigen-specific Tex at the end of all treatments (**Fig. 3d**). Transcriptional profiles of P14 cells from all groups were projected into uniform manifold approximation and projection (UMAP) space, where seven distinct scRNA-seq clusters were resolved (**Fig. 3e**). The majority of the Tex clusters resembled Tex subsets described previously ^30–32^, including: Tpex (Exh-Prog) characterized by high expression of *Slamf6* (encoding LY108); Tex-Term (Exh-Term) identified by lack of *Slamf6* and expression of *Cxcr6*, a marker of Tex terminal differentiation ^31,33^; a proliferating Tex cluster expressing cell-cycle-related genes such as *Mki67*; and a *Slamf6*-negative cluster uniquely expressing interferon-stimulated gene (ISG)-related genes (e.g. *Ifit1*) ^30,31^ (**Fig. 3e**). A small unassigned cluster was also present, but represented only ∼0.5% of all P14 cells and was not investigated further (**Fig. 3e,f**). In addition, two distinct *Slamf6*-negative “effector-like” clusters were identified by high expression of *Cx3cr1*, an Exh-Int (∼28.15% of P14) and an Exh-Int-like (∼4.94% of P14) cluster with differential expression of cell migration-related genes such as *Itga4* (**Fig. 3e,f**). The latter was mainly found in the ST^late^ treatment group (**Fig. 3g,h**).

We first used the scRNA-seq dataset to further investigate the changes of Tex subsets following different PD-1 blockade regimens. In agreement with the cytometry data (**Fig. 3a; Extended Data Fig. 4a, 5**), there was an expansion of Tex-Int and Tex-Int-like clusters in the ST^late^ and Drug Holiday treatment groups compared to Ctrl and ST^early^ treatment groups, respectively (**Fig. 3g,h**). LT PD-1 blockade resulted in an increase in the Tex-Term cluster and loss of Tex-Int cells (**Fig. 3g,h**), consistent with the cytometric analyses. These scRNA-seq data confirm that: 1) LT PD-1 blockade resulted in substantial depletion of the effector-like Tex-Int subset, the numerical erosion of Tpex subset, and enrichment of Tex-Term cells; and 2) a Drug Holiday approach was associated with the retention of the capacity to generate Tex-Int cells and preservation of the Tpex pool.

We hypothesized that prolonged PD-1 blockade in the LT group might impair the stemness of Tpex. We therefore interrogated the transcriptional signatures ^30^ specifically in Tpex from different treatment groups in the scRNA-seq data set. Consistent with the notion that PD-1 blockade promotes Tex differentiation ^3,8–10^, the transcriptional signature of Tpex biology was reduced in Tpex from the ST^late^ group, compared to the Ctrl group (**Fig. 4a**). In ST^early^ mice, however, at later time points, the canonical Tpex signature was moderately recovered (**Fig. 4a**). In contrast, LT treatment resulted in a substantial loss of the signature of Tpex biology (**Fig. 4a**), suggesting that prolonged PD-1 blockade may compromise the Tpex-associated biology in this group. To gain more insight into these Tpex changes following different anti-PD-1 treatments and the ability of Tpex to differentiate into the more the functional effector-like Tex-Int subset, we projected the scRNA-seq from only the Exh-Prog cluster of all treatment groups (in **Fig. 3d**) into a new UMAP space (**Fig. 4b**). This new UMAP projection identified five clusters of Tpex, labelled C0 to C4 (**Fig. 4b**). Gene ontology (GO) enrichment of the top differently expressed genes (DEGs) between these clusters revealed biased quiescence versus activation, inflammation, and antiviral response signatures (**Fig. 4c**). Cluster C3 was distinguished by enrichment of ISG-related transcripts, including *Ifit1*, *Ifit3*, and *Isg15* (**Fig. 4d**). In contrast, C0 had higher expression of *Tcf7*, *Myb*, and *Jun* and lacked GO term enrichment for activation or inflammatory signatures, suggesting quiescence (**Fig. 4c,d**). C0 was present in all treatment groups, with the highest proportional enrichment (∼65.07%) in the LT treatment group (**Fig. 4e**). Cluster C1 was enriched for *Klf2*, *S1pr1*, and *Cx3cr1* expression (**Fig. 4d**), and the GO terms actin cytoskeleton organization and regulation of cell migration (**Fig. 4c**), consistent with a transitioning Tpex cluster similar to the previously described Tex^prog2^ (^8^), where these cells beginning to lose progenitor biology, proliferate, and gain effector-like qualities. Although C2 and C4 clusters shared similar GO enrichment (**Fig. 4c**), cluster C2 had increased expression of genes associated with costimulation, including *Tnfrsf4* and *Icos*, whereas cluster C4 had the lowest expression of *Tcf7* but the highest in genes associated with Tex terminal differentiation (e.g. *Cxcr6* and *Entpd1*) among these Tpex populations (**Fig. 4d**). Thus, cluster C1 and C4 might represent Tpex populations poised to give rise to Tex-Int versus Tex-Term populations ^10,32^, respectively. Whereas LT treatment preferentially enriched for C0 and C4, the Drug Holiday regimen was associated with lack of C4 and strong contribution from C2 (and C1) (**Fig. 4b,e**). Altogether, these single-cell transcriptomic analyses demonstrated that LT PD-1 blockade induced transcriptional changes in Tpex, including preferential accumulation in cluster C4 that appeared poised for terminal differentiation and accumulation of cluster C0 that was enriched for stemness and quiescence.

**Fig. 4.**
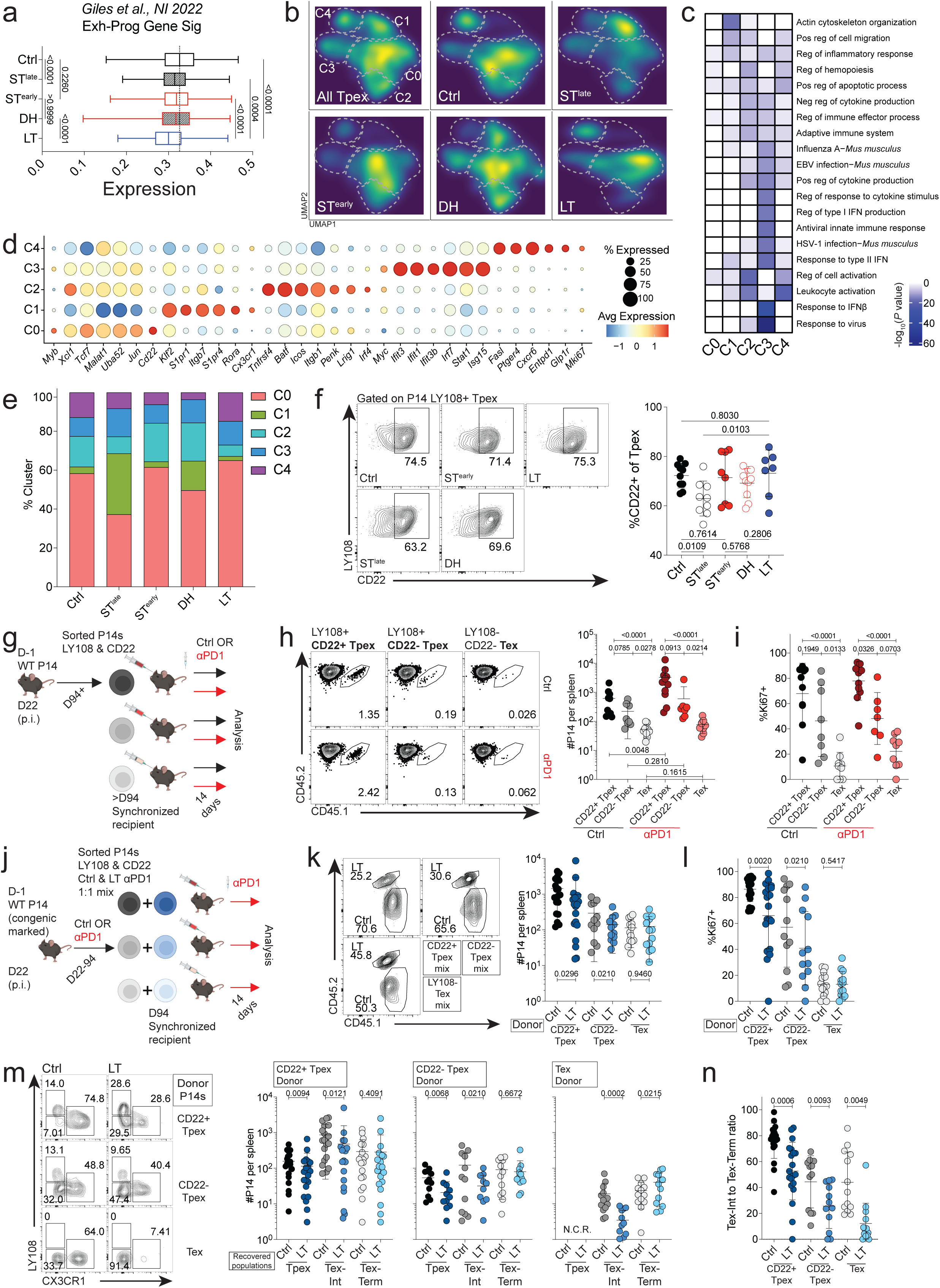
Comparison of Tpex “stemness” following different PD-1 blockade regimens. (a) Expression of Exh-Prog gene signature from ^30^ in the indicated Tpex. Dotted line shown is a reference to the median value of the Exh-Prog gene signature of Ctrl-derived Tpex. (b) UMAP of P14 Tpex following different PD-1 blockade regimens are shown. (c) Gene ontology analysis of the indicated Tpex clusters. (d) Expression of the indicated genes in the indicated Tpex cluster. (e) Proportions of different Tpex clusters summarizing (b). (f) Representative flow cytometry plots, frequencies (depicted in mean ± SD) of CD22+ cells among LY108+ Tpex in the spleen following different PD-1 blockade regimens are shown. (g) Experiment schematic of LY108 CD22 P14 Tex subset transfer experiments shown in (h and i). (h) Representative flow cytometry plots and numbers of P14 Tex (depicted in mean ± SD) recovered in spleens of recipient mice receiving the indicated donor LY108 and CD22-defined P14 Tex subsets. (i) Frequencies of Ki67+ donor P14 Tex cells. (j) Experimental schematic of LY108 and CD22-defined P14 Tex subset co-transfer experiments shown in (k-n). (k) Representative flow cytometry plots and numbers of P14 Tex (depicted in mean ± SD) recovered in spleens of recipient mice that received the indicated donor LY108 and CD22-defined P14 Tex subsets. (l) Frequencies of Ki67+ donor P14 Tex. (m) Representative flow cytometry plots and numbers of the indicated LY108 and CX3CR1-defined Tex recovered in spleens of recipient mice that received the indicated donor LY108 and CD22-defined P14 Tex subset. Note: No Cells Recovered (N.C.R.) (n) P14 Tex-Int (LY108–CX3CR1+) to Tex-Term (LY108–CX3CR1–) ratio in spleens of recipient mice that received the indicated donor LY108 and CD22-defined P14 Tex subsets. (f, h, i, k, l, m, and n) Each dot represents an individual mouse, and error bars represent SD. Statistical significance between groups was determined by (a) Dunn’s corrected Kruskal-Wallis test, (f and h) Kruskal-Wallis test, (h) two-tailed Mann-Whitney test, and (k, l, m, and n) two-tailed Wilcoxon test. Data shown in (f) (n = 7-10 per group) is representative of two independent experiments. Data shown are pooled from (h and i) two (n = 7-11 per group) or (k, l, m, and n) three (n = 12-20 per group) independent experiments.

To further interrogate the heterogeneity in the Tpex populations altered by different PD-1 blockade regimens, we examined DEGs for potential markers of Exh-Prog clusters (**Fig. 4d**). *Cd22*, an inhibitory SIGLEC family member also expressed by B cells ^34^, was expressed by cluster C0 but not by other clusters (**Fig. 4d**). CD22 protein expression was restricted to LY108+ Tpex, as predicted by the transcriptional expression (**Extended Data Fig. 8a**). Consistent with the Tpex scRNA-seq clusters (**Fig. 4e**), CD22+ Tpex were present in all treatment groups (**Fig. 4f**), although the ST^late^ treatment group had fewer CD22+ Tpex compared to Ctrl or LT mice (**Fig. 4f**), in agreement with reduced cluster C0 in this group (**Fig. 4e**).

We next investigated the kinetics the CD22+ Tpex population among P14 cells and endogenous polyclonal D^b^GP_33_tetramer+ Tex during LCMV clone 13 infection in the presence or absence of CD4 T cell help. LY108+CD22+ P14 and D^b^GP_33_tetramer+ cells were detectable by d8 of LCMV clone 13 infection and were slightly more abundant at this time point in the presence compared to the absence of CD4 T cell help (**Extended Data Fig. 8b,c**). By d15-16 p.i. the frequency of these LY108+CD22+ Tpex was similar with and without CD4 T cell help for both P14 or endogenous D^b^GP_33_-tetramer+ (**Extended Data Fig. 8b,c**). Although there were more LY108+CD22– compared to LY108+CD22+ Tpex early during chronic infection, the numerical differences normalized and there were roughly equal proportions of CD22+ and CD22– Tpex from d30-d500 p.i., though the number of both gradually declined over time during chronic antigen stimulation (**Extended Data Fig. 8d,e**). Thus, CD22 is expressed by a subpopulation of LY108+ Tpex throughout chronic infection.

*Cd22*+ Tpex had a distinct transcriptional profile, with higher expression of genes encoding IRs (e.g. *Pdcd1*, *Cd160*, and *Ctla4*) and costimulatory receptors (e.g. *Cd28* and *Cd27*) compared to *Cd22–* Tpex (**Extended Data Fig. 8f**). Transcripts for *Tcf7*, *Myb*, *Id3*, and *Eomes* – transcription factors (TFs) implicated with Tpex stemness ^5,8,27,35,36^ – were also higher in *Cd22*+ Tpex (**Extended Data Fig. 8f**). Indeed, across different PD-1 treatment groups, CD22+ Tpex consistently had higher TCF-1 expression compared to their CD22– counterparts (**Extended Data Fig. 8g**), in agreement with *Tcf7* transcript levels (**Extended Data Fig. 8f**). However, there were fewer Ki67+ cells among the CD22+ Tpex compared to CD22– Tpex in all treatment groups (**Extended Data Fig. 8h**). CD69 expression marks the more quiescent Tpex ^8^ and substantially higher proportions of CD69+ cells were found among the CD22+ Tpex (**Extended Data Fig. 8i**). *Cd22*+ Tpex also expressed some genes associated with differentiation such as *Klrd1*, *Id2*, *Gzmk,* and *Prf1*, whereas the *Cd22–* population expressed more *Tbx21, Ifng, Gzma,* and *Gzmb* (**Extended Data Fig. 8f**). A coordinated response to cellular stress may be important for Tex persistence ^30,37^ and *Cd22*+ Tpex had higher expression of stress response-related genes ^38–40^, including *Foxo1*, *Atf4*, *Xbp1, Ern1, Nfe2l1,* and *Hsf1* compared to the *Cd22–*Tpex (**Extended Data Fig. 8f**). *Cd22*+ Tpex showed selective expression of some genes associated with a previously described CD62L+ Tpex ^36^, such as *Sell*, *Myb*, *Lef1,* and *Sh3bp5*, but not others including *Bach2*, *Satb1*, *S1pr1*, or *Ccr7*, indicating only a partial overlap in biology between *Cd22*+ and CD62L+ Tpex (**Extended Data Fig. 8j**). We next investigated the relationship between the presence of CD22+LY108+ Tpex and differentiation into LY108– Tex subsets. LY108+ Tpex give rise to LY108–CX3CR1+ Tex-Int and LY108–CX3CR1– Tex-Term subsets, and this Tex subset differentiation hierarchy can be influenced by the level of persisting antigen stimulation ^8–10^. A higher fraction of CD22+ cells among Tpex was associated with a lower fraction of effector-like CX3CR1+ Tex-Int cells among the LY108– Tex (**Extended Data Fig. 8k**), suggesting that the fraction of CD22+ Tpex, or perhaps CD22 itself, may be negatively associated with the formation of the downstream Tex-Int subset. Thus, these transcriptomic and cellular analyses suggested that *Cd22*-expressing Tpex may be endowed with greater stemness potential and stress-regulating activity, while also being more restrained in their differentiation, thereby maintaining quiescence.

We next examined the proliferative and reinvigoration capacity of CD22+ versus CD22–Tpex derived from chronically infected mice. We sorted LY108+CD22+ Tpex, LY108+CD22– Tpex, and LY108–CD22– Tex P14 cells from LCMV clone 13-infected mice and adoptively transferred equal numbers of each population separately into infection-matched congenic recipient mice. These recipient mice were then left untreated or treated with anti-PD-1 for 2 weeks, and the donor Tex and their progeny were analyzed (**Fig. 4g; Extended Data Fig. 9a**). In the absence of PD-1 blockade, LY108+CD22+ Tpex resulted in the highest numerical recovery of Tex, followed by LY108+CD22– Tpex, with very few LY108–CD22– Tex progeny recovered (**Fig. 4h**). Ki67 expression was higher in the progeny of the LY108+ Tpex compared to LY108- Tex donor cells (**Fig. 4i**). Anti-PD-1-treatment resulted in reinvigoration and numerical expansion only of the donor LY108+CD22+ Tpex compared to control treatment, whereas there was limited anti-PD-1-mediated numerical expansion of LY108+CD22– Tpex or LY108–CD22– Tex (**Fig. 4h**). This anti-PD-1-mediated reinvigoration of the LY108+CD22+ Tpex was also associated with a substantially higher fraction of CD22+ Tpex donor cells expressing Ki67+ (∼80%) compared to CD22– Tpex (∼50%) or LY108–CD22– Tex (∼20%) (**Fig. 4i**), consistent with a more robust potential for expansion and reinvigoration of the CD22+ Tpex population.

These CD22+ Tpex were capable of giving rise not only to CD22– Tpex and also LY108–CD22– Tex, but also to more CD22+LY108+ cells (**Extended Data Fig. 9b**). In the absence of PD-1 blockade CD22+ Tpex gave rise to substantially more LY108+CD22+ progeny than the CD22– Tpex, indicating the self-renewing capacity of the LY108+CD22+ Tpex. In contrast, no LY108+CD22+ cells were recovered from LY108–CD22– donor cells (**Extended Data Fig. 9b**). Some CD22+ Tpex were detectable from CD22– donor Tpex suggesting some potential interconvertibility, particularly following PD-1 pathway blockade (**Extended Data Fig. 9b**). Altogether, these data indicate that CD22+ Tpex are a more stem-like Tpex population, with similarities to previous subpopulations of Tpex ^8,36^, that respond robustly to short-term PD-1 blockade.

These observations of a better response to PD-1 blockade by CD22+ Tpex made the poor responses in the setting of prolonged PD-1 blockade, where CD22+ Tpex accumulate, somewhat paradoxical. The transcriptional profile of *Cd22*-expressing Tpex revealed evidence of a heightened stress response. Together, these observations suggest that prolonged loss of PD-1 signals in LT mice may alter the biology of the CD22+ Tpex population reducing the capacity for reinvigoration. To directly compare the impact of LT PD-1 blockade on the biology of CD22+ and CD22– Tpex we performed a Tex subset co-transfer experiment using donor Tpex and Tex populations from Ctrl and LT mice treated for 10 weeks with anti-PD-1. Two groups of LCMV clone 13-infected mice received either 10 weeks of control (Ctrl) or anti-PD-1 treatment. At the end of treatment, congenically-distinct LY108+CD22+, LY108+CD22–, and LY108–CD22–Tex from Ctrl-and anti-PD-1-treated mice were purified. Equal numbers of each subset derived from Ctrl- and anti-PD-1 treated mice were mixed and adoptively transferred into infection-matched recipients of a third congenic background. These recipient mice were then treated with anti-PD-1 for 2 weeks, and the progeny of the donor P14 Tex cells was analyzed (**Fig. 4j**). Although there was no difference in reinvigoration capacity of the LY108–CD22– Tex from Ctrl versus LT anti-PD-1 mice (**Fig. 4k**), these donor cells had the lowest numerical expansion among the three donor populations (**Extended Data Fig. 9c**), as expected (**Fig. 4h**). In contrast, for both CD22+ and CD22– Tpex, the populations derived from Ctrl-treated mice outnumbered those derived from LT anti-PD-1 treated mice and there were fewer Ki67+ cells derived from either donor Tpex population from LT mice (**Fig. 4k,l**). For both donor groups, the CD22+ Tpex outperformed the CD22- Tpex (**Extended Data Fig. 9c**). Numerically, however, the CD22+ Tpex from LT anti-PD-1 treated mice gave rise to fewer LY108+CD22– Tpex and LY108–CD22– downstream Tex compared to CD22+ Tpex from Ctrl-treated mice (**Extended Data Fig. 9d**). Although CD22+ Tpex from LT mice gave rise to similar numbers of Tex-Term as CD22+ Tpex from Ctrl mice, these LT treated CD22+ Tpex gave rise to substantially fewer LY108–CX3CR1+ Tex-Int and the ratio of these more effector-like Tex-Int to Tex-Term was reduced compared to that derived from Ctrl-treated Tpex (**Fig. 4m,n; Extended Data Fig. 9d**). These data indicate that LT anti-PD-1 treatment compromises the reinvigoration potential of Tpex, particularly the CD22+ Tpex and impairs the ability of these CD22+ Tpex to give rise to the downstream effector-like Tex-Int subset.

CD22 is an ITIM-containing Siglec that binds sialic acids in *cis* or in *trans*, recruits SHP-1 and SHIP and inhibits BCR signaling in B cells ^34^. Sustained expression of CD22 by Tpex could therefore restrain Tex reinvigoration and have a role in the poor Tpex responses in LT-treated mice. We therefore next investigated whether loss of CD22 signals might overcome Tpex resistance to LT PD-1 pathway blockade. Thus, after 10 weeks of LT anti-PD-1 treatment chronically infected mice were split into a group that received another 2 weeks of anti-PD-1 or 2 weeks of anti-PD-1 plus a CD22 blocking antibody (**Fig. 5a**). These groups were compared to mice that had received no anti-PD-1 nor anti-CD22 throughout infection. Whereas anti-PD-1-only LT mice had minimal viral control compared to Ctrl mice (mainly in the liver consistent with **Fig. 1h**), addition of CD22 blockade to LT mice improved viral control in the kidney and systemically in the serum (**Fig. 5b**). To interrogate the immunological changes in these LT mice, we examined Tex in the PBMC pre- and post- the last two weeks of single versus dual treatment. The addition of anti-CD22 to LT PD-1 treatment increased the frequency and number of circulating Tex (**Fig. 5c-e**), indicating a role for CD22 blockade in Tex reinvigoration, specifically in overcoming adaptive resistance to LT PD-1 blockade.

**Fig. 5.**
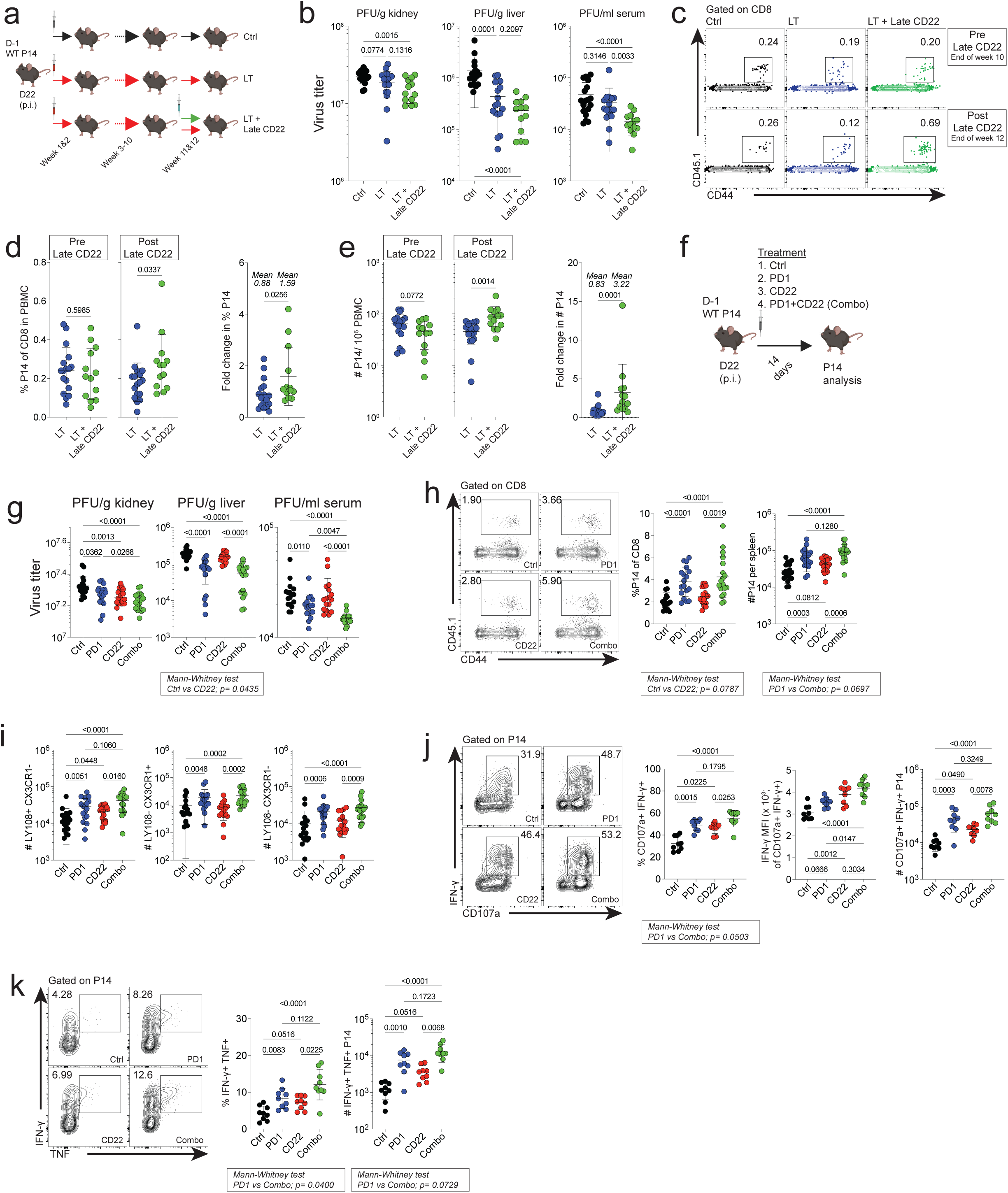
Co-blockade of CD22 and PD-1 improves Tex reinvigoration. (a) Experimental design for (b, c, d, and e). (b) LCMV viral load in kidney, liver, and serum of mice following the indicated blockade regimens are shown. (c) Representative flow cytometry plots, (d) frequencies, and (e) numbers of P14 cells in blood following the indicated blockade regimens are shown. (f) Experimental design for (g, h, i, j, and k). (g) LCMV viral load in kidney, liver, and serum of mice following the indicated blockade regimens are shown. (h) Frequencies and numbers of P14 cells following the indicated blockade regimens are shown. (i) Numbers of LY108 and CX3CR1-defined Tex subsets in the spleen following the indicated blockade regimens are shown. (j and k) Representative flow cytometry plots, frequencies and numbers of (j) CD107a and IFN-γ or (k) IFN-γ and TNF producing cells in spleen following the indicated blockade regimens are shown. (b, d, e, g, h, i, j, and k) Each dot represents an individual mouse. Statistical significance between groups was determined by (b, g, h, i, j, and k) Kruskal-Wallis test and (d, e, g, h, j, and k) two-tailed Mann-Whitney test. Data shown (b, c, d, and e) (n = 13-18 per group), (g, h, and i) (n = 17-18 per group) are pooled from two independent experiments. Data shown (j and k) (n = 9 per group) are from one independent experiment.

We next tested whether blocking CD22 improved Tex reinvigoration earlier in disease. Immunotherapy-naïve chronically infected mice were treated with anti-PD-1 and/or anti-CD22 antibodies, and viral control and Tex reinvigoration were examined (**Fig. 5f**). CD22 monotherapy resulted in modest viral control in the kidney and liver, whereas anti-PD-1 treatment resulted in substantial reduction of viral replication in the kidney, liver, and serum (**Fig. 5g**). Combination anti-CD22+anti-PD-1 further improved viral control in the kidney and serum compared to anti-PD-1 monotherapy (**Fig. 5g**). Reinvigoration of Tex was most robust for the anti-PD-1 and combination groups with trends for improved responses in the combination group in terms of number of responding Tex, number of different Tex subsets, and improved per cell function (**Fig. 5h-k; Extended Data Fig. 10a-d**). Addition of anti-CD22 to anti-PD-1 resulted in a stronger downregulation of TCF-1 in LY108+ Tpex than anti-PD-1 alone (**Extended Data Fig. 10e**). These data suggest CD22 as a target for reinvigorating Tex alone and/or in combination with PD-1.

Although the data above demonstrate CD22 expression by a subpopulation of Tpex, it was possible that effects of targeting CD22 were Tpex extrinsic given the prominent expression of CD22 by B cells ^34^. We therefore investigated whether the effect of PD-1 and CD22 co-blockade was preserved in the absence of B cells by treating chronically infected mice with the B cell-depleting anti-CD20 antibody at the time of anti-PD-1 or combination treatment (**Extended Data Fig. 11a**). Anti-CD20 resulted in a ∼98% reduction of B220+ cells indicating efficient B cell depletion (**Extended Data Fig. 11b**). Although B cell-depleted Ctrl mice showed higher viral loads in some locations than their B cell-sufficient counterparts (**Extended Data Fig. 11c**), in agreement with previous findings ^41,42^, anti-PD-1 treatment resulted in reduced viral load in untreated and B cell-depleted mice (**Extended Data Fig. 11d**). Although there was no additional benefit for viral control with combination treatment compared to anti-PD-1 alone in the kidney and liver, the enhanced control of systemic disease achieved by combined anti-CD22 and anti-PD-1 treatment was preserved in the absence of B cells with lower serum viral titers (**Extended Data Fig. 11d**). There was a consistent trend for enhanced Tex reinvigoration with combination treatment in B cell-depleted mice, although these effects did not reach statistical significance (**Extended Data Fig. 11e-g**). These data do not rule out a B cell-dependent effect of combination CD22+PD-1 co-blockade. However, B cell depletion resulted in higher viral loads in the absence of treatment (see **Extended Data Fig. 11c,d**), complicating these experiments. Moreover, loss of B cells can alter lymphoid architecture, including locations where Tpex are enriched ^6,7,41–43^ perhaps complicating the interpretation of such B cell depletion studies.

To avoid some of these challenges and further interrogate the potential of PD-1 and CD22 co-blockade to reinvigorate Tex, we engineered a mouse PD-1-CD22 bispecific antibody (BsAb). In addition, monovalent PD-1-Isotype (PD-1-Iso; i.e. one arm anti-PD-1, and one arm an irrelevant control) and CD22-Isotype (CD22-Iso) Abs were generated as controls (**Fig. 6a**). The binding characteristics of these BsAbs were evaluated using NIH-3T3 cell lines transduced to express PD-1, CD22, or both. All BsAbs demonstrated specific binding to their respective targets (**Fig. 6b-d**). The CD22-Iso Ab had improved binding compared to the PD-1-CD22 BsAb when tested against dual target-expressing cells (**Fig. 6c,d**) suggesting that addition of the monovalent anti-PD-1 arm to the monovalent anti-CD22 slightly de-tuned binding to CD22. We next tested the *in vivo* impact of these BsAbs by treating clone 13-infected mice for 2 weeks and then assessing viral control and Tex reinvigoration (**Fig. 6e**). Consistent with the data above (**Fig. 5g**), whereas PD-1-Iso or CD22-Iso each resulted in viral control in some tissues, PD-1-CD22 BsAb blockade improved viral control in all tissues compared to either mono-specific blockade and/or control treatment (**Fig. 6f**). The PD-1-CD22 BsAb also resulted in the most robust numerical reinvigoration of Tex (**Fig. 6g**) and improved the number of Tpex, Tex-Int, and Tex-Term cells compared to other treatment groups (**Extended Data Fig. 12a,b**).

**Fig. 6.**
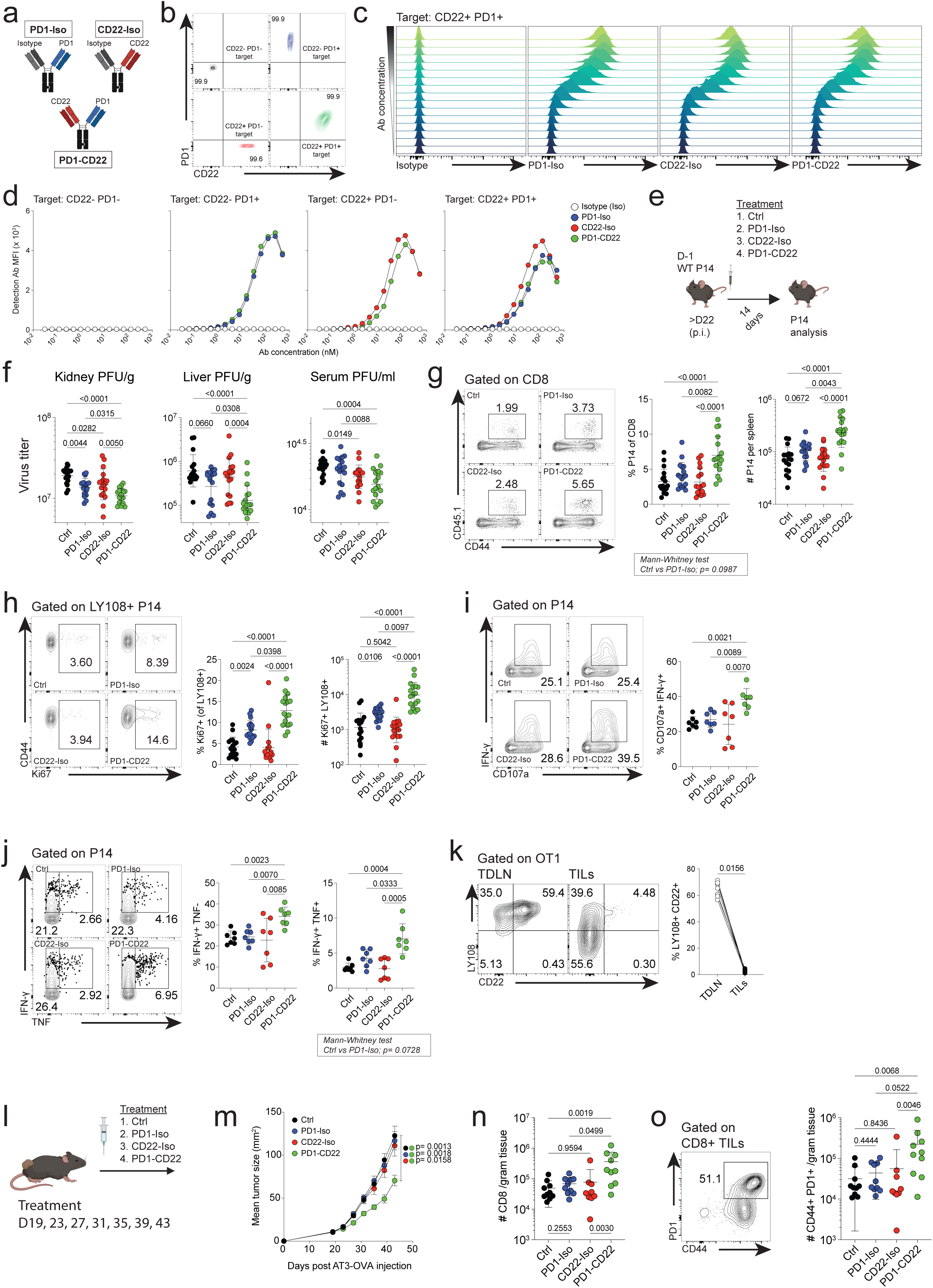
PD-1-CD22 bispecific antibody treatment reinvigorates anti-viral and anti-tumor immunity. (a) Schematic for bispecific antibodies. (b) NIH-3T3.2 cell line expressing target PD-1 and/or CD22. (c) Representative histogram plots and (d) titration summary of the indicated antibody binding to PD-1 and/or CD22. (e) Experimental design for (f, g, h, i, and j). (f) LCMV viral load in kidney, liver, and serum of mice following the indicated blockade regimens are shown. (g) Representative flow cytometry plots, frequencies, and numbers of P14 cells in spleen following the indicated blockade regimens are shown. (h) Representative flow cytometry plots, frequencies and numbers of Ki67+ cells among LY108+ Tpex in spleen following the indicated blockade regimens are shown. (i and j) Representative flow cytometry plots, frequencies of (i) CD107a+IFN-γ+ or (j) IFN-γ+TNF– and IFN-γ+TNF+ in the spleen following the indicated blockade regimens are shown. (k) Representative flow cytometry plots and frequencies of CD22+ cells among OT-1 cells in tumor draining lymph node (TDLN) and tumor-infiltrating leukocytes (TILs). (l) Experimental design for (m, n, and o). Groups of AT3-OVA-tumor bearing mice were treated with the indicated blockade regimen. (m) AT3-OVA tumor growth following the indicated blockade regimen (treatment: Day 19 to 43) are shown. Tissues were analyzed one day post last antibody treatment. (n) Numbers of intratumoral CD8+ T cells in TILs. (o) Representative flow cytometry plot and numbers of CD44+PD-1+ CD8+ TILs. (f, g, h, i, j, k, n, and o) Each dot represents an individual mouse. Statistical significance between groups was determined by (f, g, h, i, j, m, n, and o) Kruskal-Wallis test, (g, and j) two-tailed Mann-Whitney test, and (k) two-tailed Wilcoxon test. (f, g, and h) Data shown (n = 17 per group) are pooled from two independent experiments. (m) Data shown (n = 9-10 per group) are representative of two independent experiments. Data shown (i, j, and k) (n = 7 per group) and (n and o) (n = 9-10 per group) are from one independent experiment.

Treatment with the PD-1-CD22 BsAb also resulted in robust increases in the frequency and number of proliferating (Ki67+) Tpex compared to Ctrl or either mono-specific Ab (**Fig. 6h**). In contrast, B cell numbers and proliferation remained unchanged in all treatment groups (**Extended Data Fig. 12c,d**), consistent with the observations in mice treated with separate anti-PD-1 and anti-CD22 antibodies (**Extended Data Fig. 11b**). In addition to Tex numerical expansion, PD-1-CD22 BsAb treatment also resulted in improved effector function with increased cytokine production, degranulation, and the fraction of Tex co-producing effector cytokines (e.g. IFN-γ and TNF) (**Fig. 6i,j**). These data demonstrate the combination targeting of PD-1 and CD22 using a BsAb robustly reinvigorated Tex with improved viral control, Tex numerical expansion, and increased per cell function compared to single targeting of PD-1 or CD22 alone.

We next investigated whether CD22 had a role in tumor-specific Tpex and whether PD-1-CD22 BsAb treatment could improve anti-tumor immunity. LY108+ (or TCF-1+) Tpex in tumor-draining lymph node (TDLN) and/or tumors have a key role in anti-tumor immunity ^44–47^. Mice containing OT-1 cells were implanted with AT3-OVA tumors and on d32 post tumor inoculation the donor OT-1 cells in the TDLN and tumor were examined. In the tumor, OT-1 cells were largely LY108– Tex and there were few LY108+CD22+ TIL (**Fig. 6k**). However, the tumor-specific OT-1 cells in the TDLN were largely LY108+, with low expression of IRs such as PD-1 and LAG3 (**Extended Data Fig. 12e**), consistent with previous work ^44–47^. However, in contrast to TIL, a larger proportion of tumor-specific CD22+ Tpex was found in the TDLN (**Fig. 6k**). We next tested treatment with the PD-1-CD22 BsAb in the AT3-OVA tumor model without OT-1 cells to assess endogenous anti-tumor immunity (**Fig. 6l**). In this setting, treatment with the monovalent anti-PD-1-Iso or anti-CD22-Iso Abs had no impact on tumor growth compared to isotype-treated (**Fig. 6m**). Treatment with the PD-1-CD22 BsAb, however, resulted in a reduced tumor growth (**Fig. 6m**) and was associated with a robust expansion of intratumoral total and PD-1+ CD8 T cells (**Fig. 6n,o**). Thus, co-targeting PD-1 and CD22 using a BsAb had benefits in both chronic viral infection and cancer. Overall, these data identify a role for CD22 expression on Tpex in coregulating these progenitor cells in multiple disease contexts. Moreover, these observations indicate that a failure to sustain Tex reinvigoration during long-term treatment with anti-PD-1 may be overcome by targeting CD22 for improved reinvigoration.

The immunopharmacology of PD-1 pathway targeting drugs remains poorly understood. Although the pharmacodynamics of the antibodies used to block this pathway are well documented ^25^, the “pharmacology” of the reinvigorated Tex cells that are the major *in vivo* effector mechanism of anti-PD-1 induced physiological changes has remained less well defined. Preclinical studies have been limited by a paucity of models where tumors do not grow progressively, complicating logistics and interpretation of long-term treatment approaches. Moreover, most previous preclinical studies have used species mismatched blocking antibodies (i.e. rat IgG) limiting long-term or retreatment strategies due to the induction of anti-rat antibodies. Finally, precisely tracking responding CD8 T cells of known specificity and stimulation history is often challenging in both preclinical cancer models and in humans. Here, we employed the chronic LCMV infection model with longitudinally trackable antigen-specific CD8 T cells, and mouse anti-mouse PD-1 blocking antibodies to interrogate these questions. Our data demonstrate that an intermittent “Drug Holiday” PD-1 blockade strategy permits a secondary round of Tex reinvigoration that is not observed during continuous long-term PD-1 pathway blockade. This secondary reinvigoration is comparable to the Tex reinvigorated observed during an initial PD-1 blockade in both T cell dynamics and control of disease burden. The finding that prolonged continuous PD-1 blockade had little benefit on sustaining or increasing Tex reinvigoration after the initial burst of reinvigoration is consistent with a single peak of immune reinvigoration observed in human melanoma patients ^11,12^ and Tex-intrinsic adaptive resistance to long-term loss of PD-1 signals.

Mechanistically, the different PD-1 blockade treatment regimens rebalanced subsets of Tex with potentially therapeutically relevant increases in the effector-like Tex-Int population in the Drug Holiday but not continuous (LT) PD-1 blockade. Moreover, depletion of the effector-like Tex-Int population, changes in PD-1 expression on Tpex populations with continuous PD-1 pathway blockade, and/or co-expression of other IRs with long-term anti-PD-1 treatment may have implications for clinical combination treatment strategies. The approach used here included analysis of both polyclonal and TCR transgenic CD8 T cells allowing us, in the latter case, to exclude contributions from new T cell priming. It is possible in settings where T cell priming can occur, continuous PD-1 blockade may have additional effects. However, our data highlight the potential value of providing periods of “rest” from PD-1 pathway blockade followed by cycles of retreatment.

The dynamics of Tex reinvigoration, which varied across different PD-1 blockade regimens, suggest a reassessment of the optimal duration and scheduling PD-1 pathway-targeted therapy. Current clinical PD-1 blocking reagents have been optimized for high affinities and slow off rates resulting in some cases in receptor occupancy of >70% even when serum concentrations are undetectable and a receptor occupancy half-life of several months after a single infusion ^25^. These drug features might complicate strategies aimed at providing a drug holiday in humans. Nevertheless, several clinical observations support the potential benefit of retreatment after a washout of anti-PD-1 (or anti-PD-L1). For example, in small studies clinical responses were observed in 14.7% (5/34 patients) to 31.5% (6/19 patients) of melanoma patients that had undergone anti-PD-1 retreatment due to disease progression after a treatment-free period of several weeks to >18 months ^48,49^. Although the benefit of removing a patient from long-term PD-1 pathway blockade therapy may vary depending on the disease type and stage ^48–56^, as well as the presence of immune-related adverse events, these clinical observations support the notion that a drug holiday may have utility. Such strategies might have benefit for limiting immune related adverse events and for combination therapies where new therapeutics could be added in subsequent cycles of PD-1 blockade to capitalize on the burst of Tex reinvigoration that occurs upon retreatment.

CD22+ Tpex were generated early during chronic infection and were maintained long-term during chronic infection similarly to their CD22– Tpex counterparts. Although heterogeneity in the Tpex compartment has been previously observed ^8,36^, CD22 may confer a biological advantage to CD22+ Tpex under certain conditions. During chronic infections, the *raison d’être* for forming a pool of Tex may be to establish a host–pathogen stalemate allowing the host to survive through reproductive age ^29^. To establish such a balance chronically stimulated Tex tune TCR signaling threshold, in part using PD-1, to establish this homeostasis ^57–60^. Indeed, if PD-1 is permanently removed genetically, Tex can form, but the population erodes over time leading to loss of Tex long-term ^28,61,62^. Moreover, PD-1 signaling is important for the formation and/or preservation of key Tpex-like cells in acute infection ^63^. In addition, PD-1 appears to be essential for Tpex cells to balance self-renewal and proliferation in settings of strong antigen stimulation ^64^. Long-term PD-1 pathway blockade may also result in a similar excessive Tex stimulation, proliferation, and differentiation into downstream Tex subsets resulting in greater dependency on other negative regulation and/or the induction of cell stress response pathways. Such a notion is consistent with recent studies identifying proteostatic stress as a key adaptation in Tex ^65,66^. Moreover, *Cd22*-expressing Tpex also expressed several TFs (e.g., *Foxo1*, *Atf4*, *Xbp1, Ern1, Nfe2l1,* and *Hsf1*) involved in regulating endoplasmic reticulum (ER) stress from misfolded proteins, oxidative stress, and heat shock response ^38–40^. We speculate that these stress-regulating mechanisms may help preserve the CD22+ Tpex by limiting proliferation and differentiation. Following long-term PD-1 blockade, the remaining Tpex were skewed in their ability to give rise to downstream Tex subsets with a relative loss in the ability to form the more cytotoxic effector-like Tex-Int subset, instead giving rise to more terminal Tex. Such a shift may limit tissue pathology from the Tex-Int subset in some settings, but results in less optimal disease control in the setting of chronic infections and cancer.

Chronic stress, injury, or aging are often associated with dysfunction of tissue stem and progenitor cells ^67–69^. Tissue stem and progenitor cells often exist as two related populations: a more active, cycling population that maintains tissue homeostasis during normal conditions, and a reserve, typically quiescent population that can be reactivated during tissue damage or stress ^67–71^. After acutely resolved stress, the quiescent tissue stem cells that were mobilized into proliferation and differentiation to help regenerate the tissue, then return to quiescence to maintain future responsiveness ^70–73^. In contrast, chronic stress can erode or “exhaust” the hematopoietic or other tissue stem cell compartments, leading to defects in long-term repopulation and/or differentiation ^74–76^.

There may be analogies to these developmental biology frameworks for Tex. Indeed, Tpex have the ability to self-renew and give rise to several epigenetically distinct differentiated downstream Tex subsets during both homeostatic conditions and when reinvigorated by checkpoint blockade ^30,31^. Moreover, different subpopulations of Tpex have been described ^8,36,77^, including those that are more active and more quiescent ^8^. One possible consequence of long-term PD-1 pathway blockade is to force excessive proliferative and differentiation stress on Tpex, eroding the ability of this progenitor cell compartment to continually give rise to downstream progeny. In contrast, Tpex that had undergone a drug holiday, providing some relative rest from excessive proliferative and differentiation stress, regained their ability for secondary anti-PD-1 mediated Tex reinvigoration. Co-expression of IR like CD22, (and perhaps others like LAG3 ^8,9,63^) with PD-1 by Tpex contrasts with other IRs, such as Tim3 and Tigit that are preferentially expressed by more terminal Tex ^7–10,20,36,78–81^. CD22 potentially through SHP-1/2 or SHIP mediated attenuation of TCR signaling may provide an additional mechanism to restrain over stimulation of Tpex and counteract erosion of this key progenitor population over time. While better known for its role on B cells, CD22 expression has been reported on a population of CD8 T cells during helminth infection where this IR regulates the CD8 T cell effector program ^82^. Moreover, CD22+ Tregs may have a role in <u>M</u>ultisystem Inflammatory <u>S</u>yndrome in <u>C</u>hildren (MIS-C) associated with SARS-CoV-2 infection in humans ^83^. Thus, co-targeting of IRs, like CD22, expressed preferentially by Tpex may provide advantages for immune reinvigoration. Moreover, the enrichment of Tpex subject to additional negative regulatory restraints due to long-term PD-1 pathway blockade points to targetable mechanisms to overcome Tex-intrinsic adaptive resistance to immunotherapy through two new concepts: (1) targeting Tpex-expressed stress-associated negative regulatory pathways; and (2) cycles of treatment, rest and retreatment with checkpoint blockade immunotherapies.

## Materials and Methods

### Mice

C57BL/6 mice were purchased from Charles River (NCI) or Jackson Laboratories. Wild type (WT) C57BL/6 congenically distinct P14 (T cell receptor transgenic specific for GP^33–41^ in CD8 T cells) mice were bred as described ^84,85^. WT C57BL/6 congenically distinct OT-1 (T cell receptor transgenic specific for OVA^257–264^ in CD8 T cells) mice were bred as described ^86,87^. Mice were maintained in SPF facility at the University of Pennsylvania (UPenn). Experiments and procedures were performed in accordance with approval from the UPenn Institutional Animal Care and Use Committee (IACUC).

### Adoptive transfer of P14 cells

P14 cells were isolated from the peripheral blood of donor mice using gradient centrifugation (Histopaque-1083; Sigma-Aldrich). A total of 500 to 1,000 congenically marked P14 cells were adoptively transferred i.v. into 5-6 week old C57BL/6 mice one day prior to infection. This number of P14 cells (500 to 1,000 cells) does not impact LCMV clone 13 viral load ^21,28^.

### LCMV clone 13 infection and antibody treatment

LCMV clone 13 was propagated and titers were determined as described ^19,28^. C57BL/6 mice containing adoptively transferred P14 cells were infected i.v. with 4 x 10^6^ plaque forming units (PFU) of LCMV clone 13. Unless otherwise indicated, CD4 T cells were transiently depleted by i.p. injection of 200 𝜇g anti-CD4 depleting antibody (clone GK1.5; Bio X Cell) on days -1 and +1 p.i. Anti-PD-1 (clone RMP1-14, mouse IgG1/IgG2a-D265A, 200 𝜇g/injection; Absolute Antibody, Bio X Cell, or Leinco Technologies) or control was injected i.p. every 3 days starting as indicated in experimental schematics (**Fig. 1a, 2a, 4g, 4j, 5a,** and **5f**). Anti-PD-1 (clone RMP1-14, rat IgG2a, 200 𝜇g/injection; Bio X Cell) or control was injected i.p. every 3 days starting from d22 p.i., as indicated in experimental schematics (**Extended Data Fig. 7a**). Anti-CD22 (clone Cy34.1, mouse IgG1, 200 𝜇g/injection; Bio X Cell) was injected i.p. every 3 days starting as indicated in experimental schematics (**Fig. 5a** and **5f**). Anti-PD-1/CD22 bispecific (mouse IgG1-D265A, 100 𝜇g/injection; see below) was injected i.p. every 3 days starting as indicated in experimental schematics (**Fig. 6e**). Anti-CD20 (clone MB2-011, mouse IgG2c, 200 𝜇/injection; Bio X Cell) was injected i.p. every 3 days starting as indicated in experimental schematics (**Extended Data Fig. 11a**). Viral load for the indicated organs was determined as described ^19,20^.

### Bispecific antibody design, synthesis, and binding validation

To create the anti-PD-1/CD22 bispecific antibody, heavy and light chain sequences were sourced from anti-PD-1 (clone RMP1-14) and anti-CD22 (clone Cy34.1) and engineered into a mouse IgG1 backbone containing a D265A modification, using a “knob-in-hole” design and the mouse version of CrossMab approaches for forced chain pairing ^88–90^. The antibody was synthesized using Biointron services. The binding specificity of anti-PD-1/CD22 was validated using PD-1- and/or CD22-expressing target cell lines (**Fig. 6b-d**). In brief, an unconjugated anti-PD-1/CD22 bispecific was used to stain target cell lines. Secondary fluorochrome-conjugated anti-mouse IgG1 was used to detect target cell-bound bispecific antibody.

### Mouse tumor models and antibody treatment

MCA1956 (1 x 10^6^ cells; kindly provided by Dr. Robert Schreiber, Washington University, St. Louis) or AT3-OVA (2 x 10^5^ cells; kindly provided by Dr. Paul Beavis, Peter MacCallum Cancer Centre) tumor cells were injected subcutaneously into C57BL/6 mice. For the AT3-OVA model, naïve OT-1 cells (1 x 10^5^ cells) were adoptively transferred into tumor-bearing mice on d6 post tumor inoculation. Anti-PD-1 (clone RMP1-14, mouse IgG2a-D265A, 200 𝜇g/injection; Bio X Cell) or control was injected i.p. every 4 days starting as indicated in experimental schematics (**Extended Data Fig. 2a**). The anti-PD-1/CD22 bispecific (mouse IgG1-D265A, 100 𝜇g/injection; in-house) was injected i.p. every 4 days starting as indicated in experimental schematics (**Fig. 6l**). Tumor size was monitored using a digital caliper, and tumor area (length x width; mm^2^) was determined.

### Sample processing

For longitudinal monitoring of responses, peripheral blood mononuclear cells were isolated using gradient centrifugation (Histopaque-1083; Sigma-Aldrich), at the indicated time points shown. For liver analysis, the gall bladder was removed, and liver tissue was homogenized. Cells were resuspended in 5 ml of 40% Percoll solution (GE Healthcare) and underlaid with 2 ml of 60% Percoll solution. Cells were spun at 2000 rpm for 20 min at 20°C, and the lymphocyte population was collected from the interface. For spleen, pieces of spleen mice were homogenized and passed through a 70 𝜇m cell strainer. Red blood cells were lysed in ACK Lysis Buffer (Gibco; Thermo Fisher Scientific), then washed in PBS supplemented with 1% FBS, before passing through a 70 𝜇m cell strainer. For in vitro restimulation, splenocytes were incubated for 5 hours at 37°C in the presence of GolgiPlug (BD) and GolgiStop (BD) with GP^33–41^ peptide (KAVYNFATM; final 0.2 𝜇g/ml)_­._

For tumor experiments, tissue analysis was performed on tumor draining lymph nodes (TDLN; inguinal lymph node of tumor injection side), and tumors (as tumor-infiltrating lymphocytes, TILs) at the indicated time points. Lymph nodes were processed through a 70 𝜇m cell strainer. For TILs, minced tumor tissues were incubated in digestion media [DMEM with Type 4 Collagenase (1 mg/mL) and DNaseI (0.02 mg/mL)] at 37°C for 40 min, and processed through a 70 𝜇m cell strainer. BD Liquid Counting Beads were used to quantify the number of TILs ^85^.

### Flow cytometry

Single cell suspensions were stained with Live/Dead Dye. Surface staining was then performed at 4°C for 30 min. When applicable, secondary staining using fluorochrome-conjugated Streptavidin was performed at 4°C for 30 min. Intracellular staining of transcription factors, Ki67, and Granzyme B was performed using the eBioscience Foxp3/Transcription Factor Staining Buffer Set according to the manufacturer’s protocol. Intracellular staining for IFN-y, TNF, and Mip1a was performed using the BD Cytofix/Cytoperm kit according to the manufacturer’s protocol. Live cells were discriminated using Live/Dead dye. Stained samples were acquired on BD LSR II and Symphony A5 machines. Voltages on the machine were standardized using Spherotech rainbow beads (SRCP-35-5A and RFP-30-5A). Flow cytometry data were processed and analyzed using FlowJo software (TreeStar).

### Tex subset sorting and transfer experiments

P14 cells from the indicated time points were enriched using CD8 T cell negative isolation kit (EasySep Mouse CD8 T cell isolation kit Cat#19853, STEMCELL Technologies) and stained for the donor congenic marker, Live/Dead Aqua dye, LY108, and CD22. Sorted LY108 and CD22 defined Tex subsets (9 x 10^3^ to 1 x 10^4^ cells, each subset) were adoptively transferred into congenically-distinct recipient mice with synchronized infections. Anti-PD-1 (clone RMP1-14, mouse IgG2a-D265A, 200 𝜇g/injection; Leinco Technologies) or control antibody was injected i.p. every 3 days starting as indicated in experimental schematics (**Fig. 4g** and **4j**).

### scRNA-seq library generation

The scRNA-seq library was generated using the 10x Genomics Chromium Single Cell 5’ v2 (Dual Index) Library Kit. In brief, sorted P14 cells from different PD-1 blockade regimens were washed with 0.04% BSA PBS, and ∼3,700 to 25,000 cells were loaded into a 10x Chromium Controller (Chromium Next GEM Chip K). Library preparation was performed according to the manufacturer’s guidebook. Libraries were checked using an Agilent Tapestation, then quantified using a KAPA Library Quantification Kit and sequenced on an Illumina NovaSeq.

### scRNA-seq data processing and analysis

scRNA-seq data were generated using the 10x Cell Ranger pipeline (6.0.0) and mm10 genome. We generated fastq files using cellranger mkfastq, quantified reads using cellranger count, and cellranger aggr to combine samples. Downstream analysis was performed in R (version 4.4.1) and Seurat (version 4.0.5 and 5.2.0) using default parameters unless otherwise noted. Cells with less than 200 features (but no more than 3500 features), and more than 5% mitochondrial reads were excluded. Standard Seurat data processing and normalization steps were performed: RunPCA, RunUMAP, FindNeighbors and FindClusters; clusters with low-quality scores were removed, and the final resolution was 0.2 for all P14 (**Fig. 3e**), and 0.25 for Tpex (**Fig. 4b**) UMAPs, respectively. The gene expression signature of Exh-Prog (**Fig. 4a**) was from Giles et al ^30^. DEGs of Tpex clusters C0-C4 (**Fig. 4d**) were generated using default FindMarkers function (Seurat) with a log2 fold change threshold of 0, and filtered with a false discovery rate (FDR) of less than 0.05 using the Benjamini-Hochberg method to adjust p values. Gene set enrichment was performed using the AddModuleScore function (Seurat). DotPlot (Seurat) was used to illustrate the indicated genes for the indicated populations (**Fig. 4d**; and **Extended Data Fig. 8f,** and **8j**). Gene Ontology (GO) analysis of DEGs was performed using Metascape (https://metascape.org/) (**Fig. 4c**).

### Statistical analysis

Statistical tests for data were performed using GraphPad Prism software. Where applicable, outliers were excluded using GraphPad Prism’s outlier removal (ROUT) method with *Q* set at 1%. A two-tailed Mann-Whitney test was used for comparisons between two treatment groups. A two-tailed Wilcoxon test was used for comparisons between paired samples. A Kruskal-Wallis test was used for comparisons between more than two treatment groups. A Log-rank test was used to assess survival differences between treatment groups. Correlation analysis between two variables was assessed using Pearson correlation coefficient. A *p* value of <0.05 was considered significant in these analyses (* p < 0.05; ** p < 0.01; *** p < 0.001; **** p < 0.0001; ns, not significant; or numerical p values as indicated in the graph).

## Acknowledgements

We thank members of the Wherry laboratory for helpful discussions and critical analysis of the manuscript. We also thank A.C. Huang for helpful discussions. We thank the Penn Cytomics and Cell Sorting Shared Resource Laboratory for providing technical support and instrumentation. We thank V. Ekshyyan, C.W. Lau, M. McLaughlin, C.H. Holliday, D. Morrall-Anderson, and B. DeGregorio for laboratory support. We thank the NIH Tetramer Core Facility for tetramer reagents.

## Contributions

Conceptualization: SFN, EJW

Methodology: SFN, EJW, SM, JRG, DM

Investigation: SFN, RPO, YJH, JRG, VA, DM, MK, KN, ZC, AEB, KPP, JEW, RPS, MAA

Visualization: SFN, SM, ARG, DM, EJW

Funding acquisition: SFN, EJW

Project administration: SFN, EJW

Supervision: SFN, EJW, DW

Writing – original draft: SFN, EJW

Writing – review & editing: SFN, SM, RPO, YJH, JRG, VA, DM, MK, KN, ZC, AEB, KPP, JEW, RPS, MAA, ARG, DW, EJW

## Data and materials availability

All data are available in the main text or the supplementary materials.

---

**Extended Data Fig. 1.**
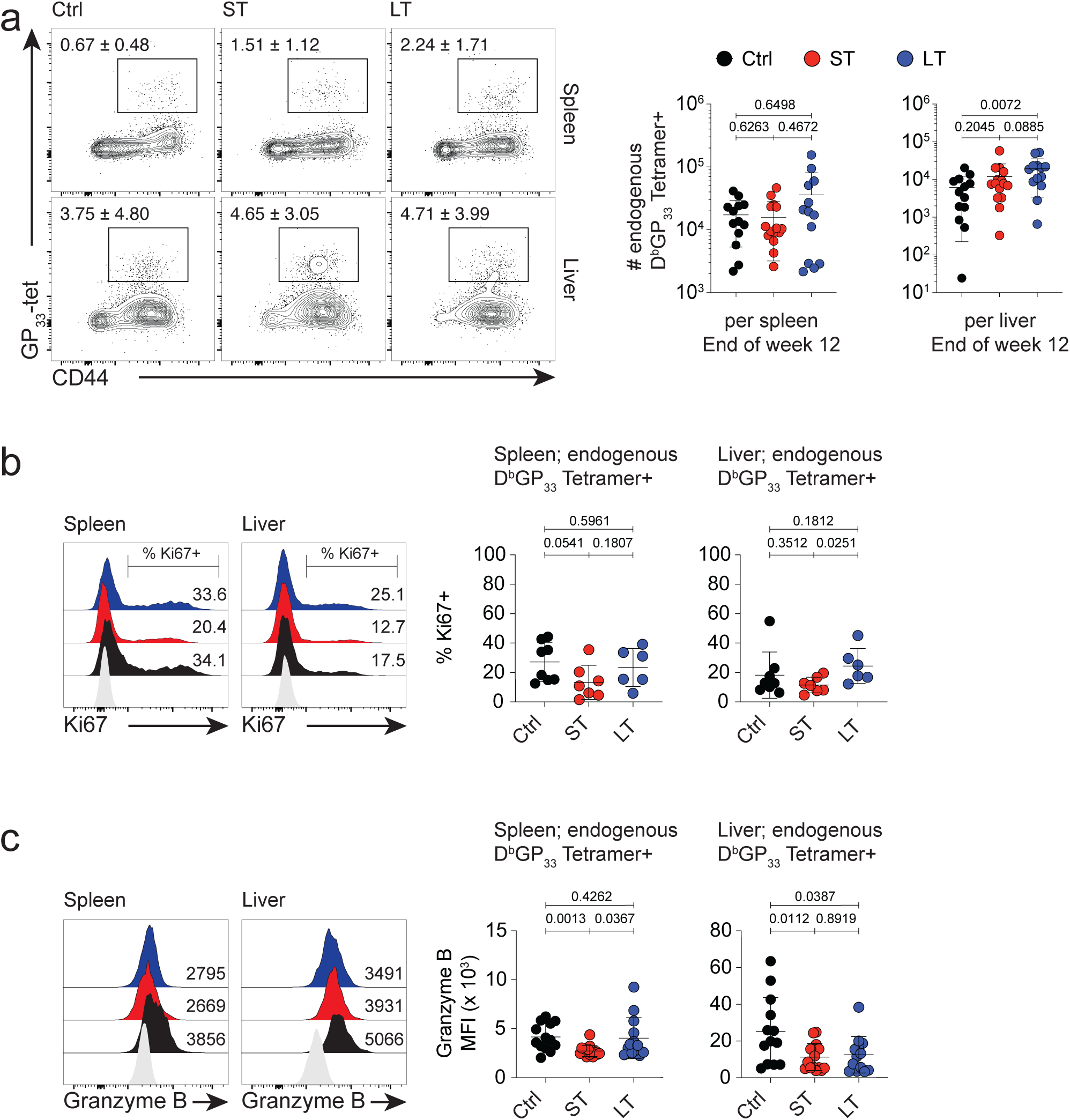
Comparison of antigen-specific Tex responses between transient and long-term PD-1 blockade. Groups of LCMV clone 13-infected B6 mice containing adoptively transferred P14 cells were treated with the indicated PD-1 blockade regimen as Fig. 1a. (a) Number of endogenous D^b^GP_33_-tetramer+ CD8 T cells in the spleen and liver are shown. (b) Frequencies of Ki67+ cells as a fraction of endogenous D^b^GP_33_-tetramer+ CD8 T cells are shown. (c) MFI for Granzyme B expression by endogenous D^b^GP_33_-tetramer+ CD8 T cells are shown. Each dot in (a), (b), and (c) represents an individual mouse, and error bars represent SD. Statistical significance between groups was determined by two-tailed Mann-Whitney test. Data shown in (a) and (c) (n = 13-15 per group) are pooled from two independent experiments. Data shown in (b) (n = 6-8 per group) are from one independent experiment.

**Extended Data Fig. 2.**
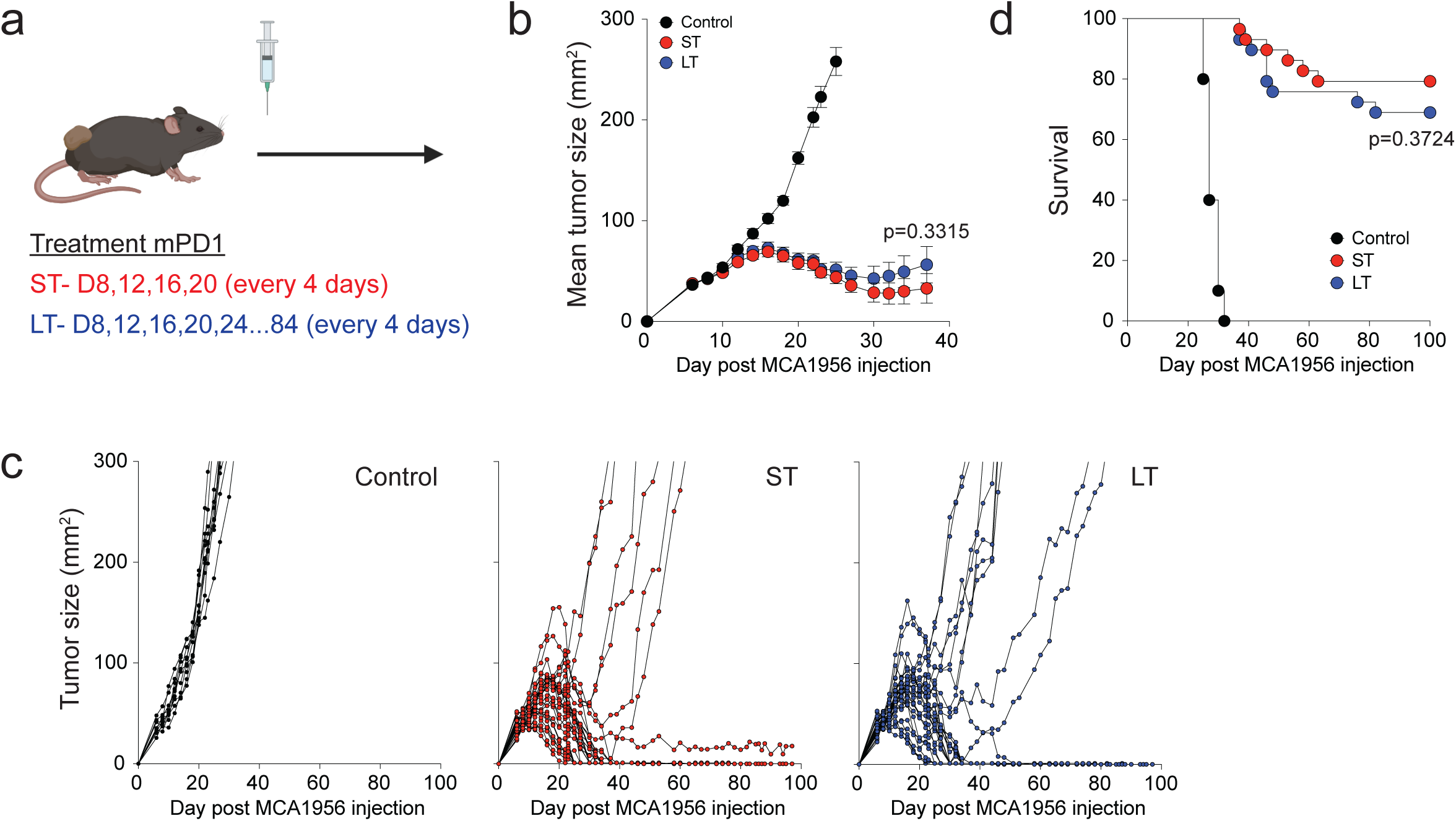
Short-term and long-term PD-1 blockade in a mouse tumor model. (a) Groups of MCA1956-tumor bearing mice were treated with PD-1 blockade regimens as shown. (b) Mean tumor growth curve (mean ± SEM), (c) overall survival, and (d) individual tumor growth of individual mice are shown. Each line in (c) represents an individual mouse (n = 10-29 per group). Statistical significance between ST versus LT determined by (b) two-tailed Mann-Whitney test and (d) Log-rank test.

**Extended Data Fig. 3.**
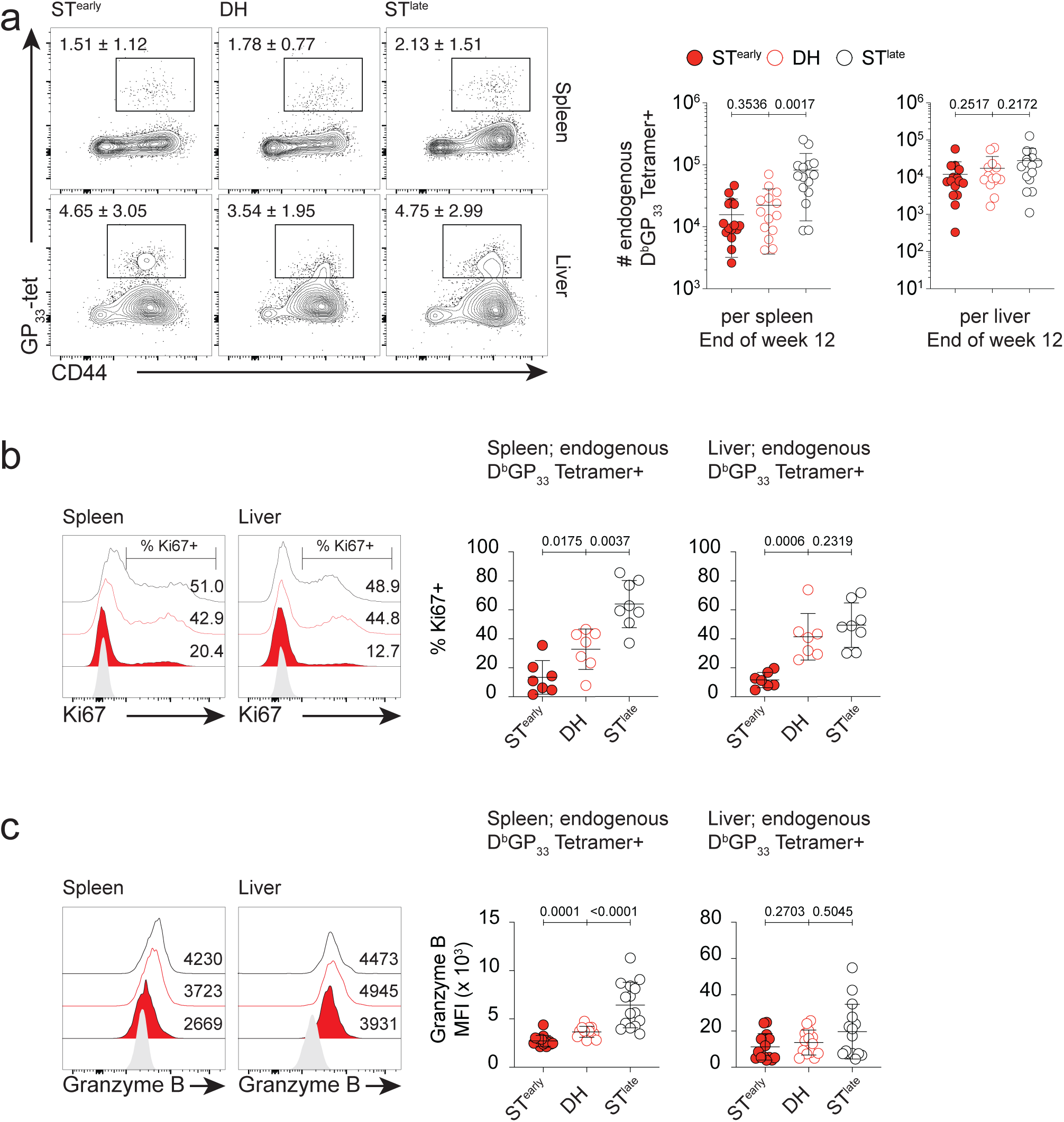
Comparison of antigen-specific Tex responses from ST^early^, Drug Holiday, and ST^late^ treatment groups. Groups of LCMV clone 13-infected B6 mice containing adoptively transferred P14 cells were treated with the indicated PD-1 blockade regimen as Fig. 2a. (a) Numbers of endogenous D^b^GP_33_-tetramer+ CD8 T cells in the spleen and liver are shown. (b) Frequencies of Ki67+ cells as a fraction of endogenous D^b^GP_33_-tetramer+ CD8 T cells are shown. (c) MFI for Granzyme B expression by endogenous D^b^GP_33_-tetramer+ CD8 T cells are shown. Each dot in (a), (b), and (c) represents an individual mouse, and error bars represent SD. Statistical significance between groups was determined by two-tailed Mann-Whitney test. Data shown in (a) and (c) (n = 14-15 per group) are pooled from two independent experiments. Data shown in (b) (n = 7-8 per group) are from one independent experiment.

**Extended Data Fig. 4.**
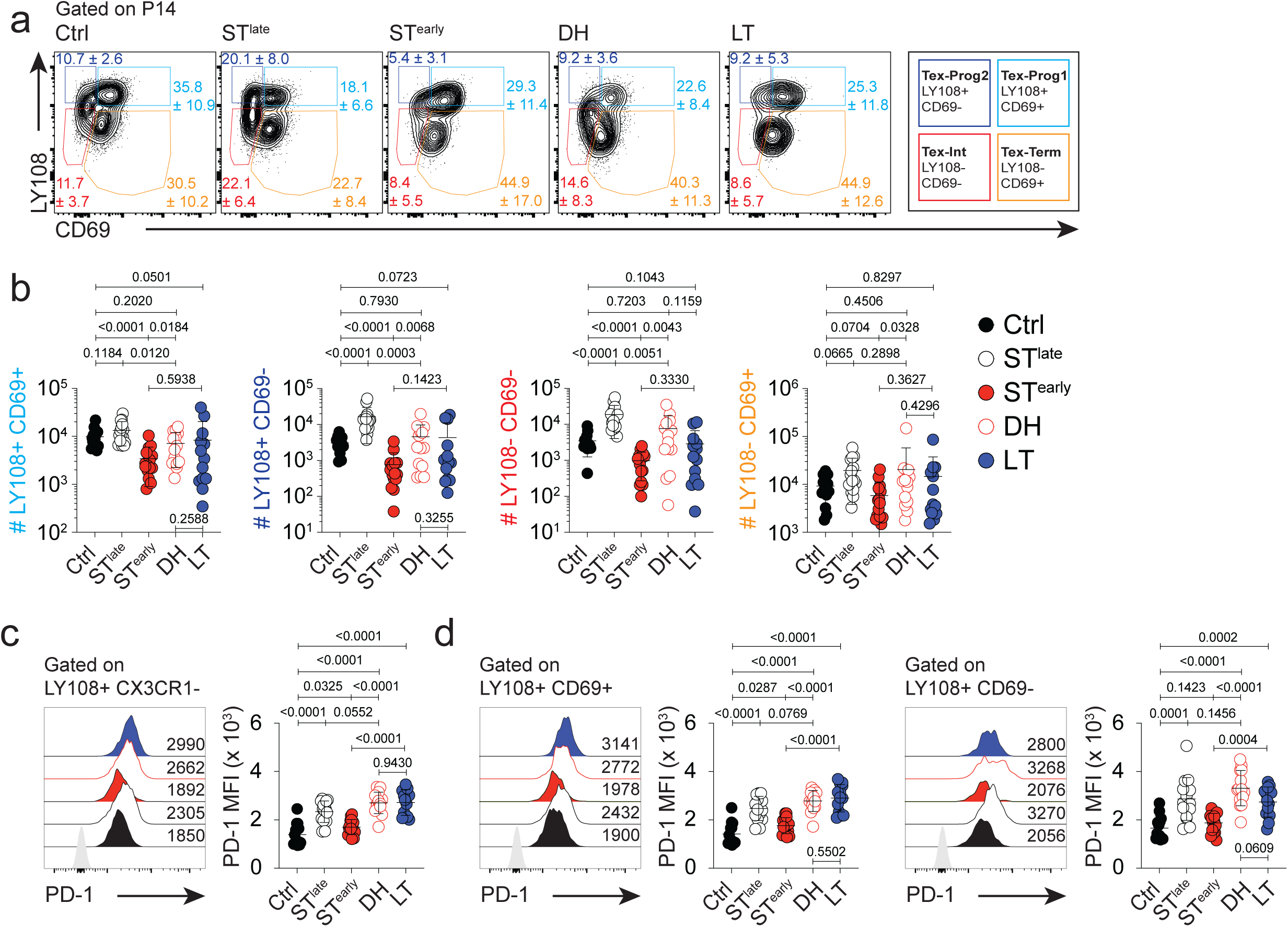
Comparison of P14 Tex responses following different PD-1 blockade regimens. (a) Representative flow cytometry plots, frequencies (depicted in mean ± SD; See Extended Data Fig. 5b) and (b) numbers of LY108 and CD69-defined Tex subsets in the spleen following different PD-1 blockade regimens are shown. Representative overlaid histograms and MFI of PD-1 expression for (c) LY108+CX3CR1–, (d) LY108+CD69+ (left panel) and LY108+CD69– (right panel) P14 cells are shown. Grey histogram in (c) and (d): gated on CD44– naïve CD8 T cells. (b, c, and d) Each dot represents an individual mouse, and error bars represent SD. Statistical significance between groups was determined by two-tailed Mann-Whitney test. (a, b, c, and d) Data shown (n = 13-15 per group) are pooled from two independent experiments.

**Extended Data Fig. 5.**
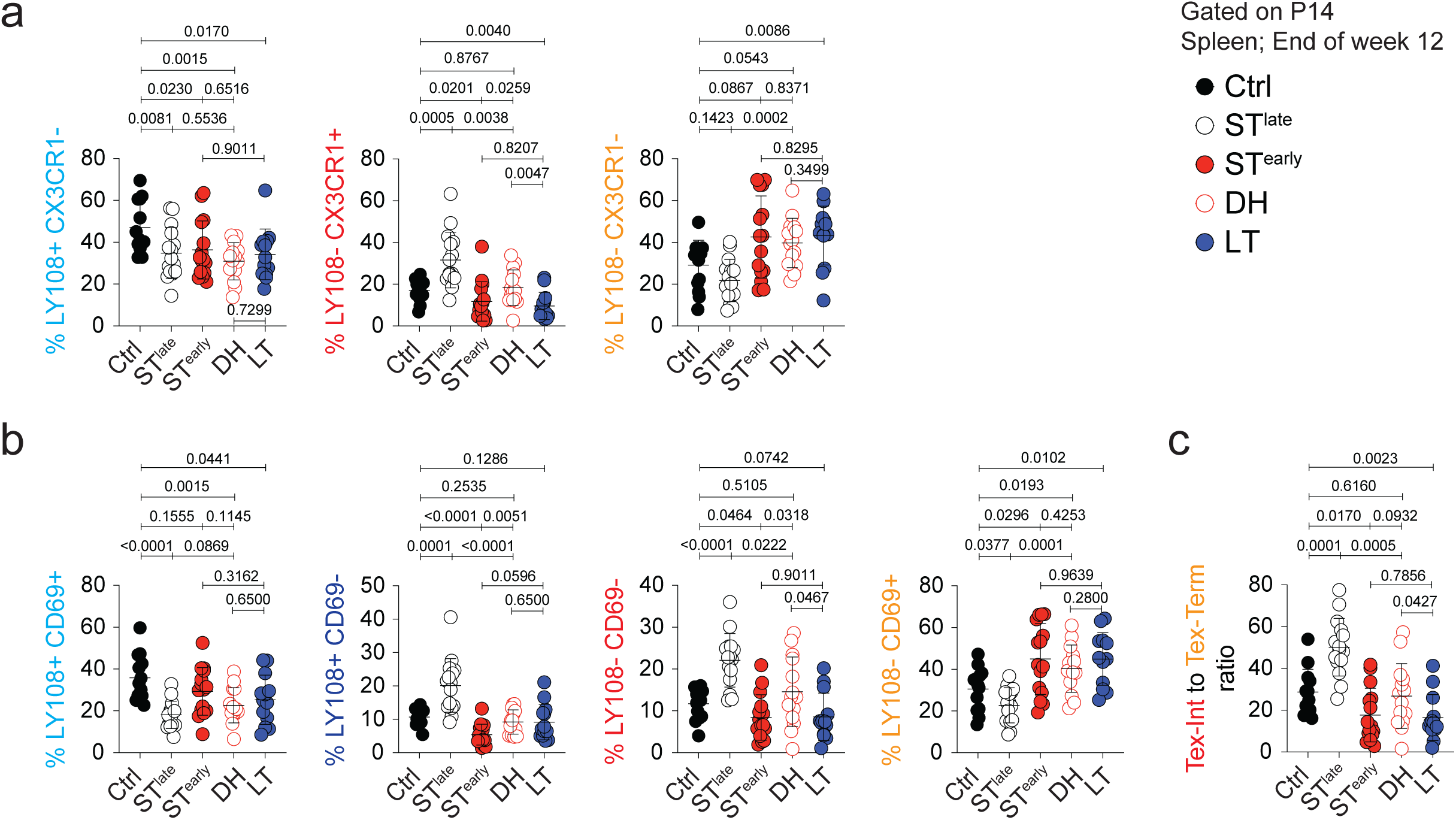
Tex subset distribution for P14 cells following different PD-1 blockade regimens. Frequencies of (a) LY108 CX3CR1 and (b) LY108 CD69 Tex subsets for P14 cells following different PD-1 blockade regimens are shown. (c) P14 Tex-Int (LY108–CD69–) to Tex-Term (LY108–CD69+) ratios in spleen following different PD-1 blockade regimens are shown. Each dot represents an individual mouse, and error bars represent SD. Statistical significance between groups was determined by two-tailed Mann-Whitney test. (a, b, and c) Data shown (n = 13-15 per group) are pooled from two independent experiments.

**Extended Data Fig. 6.**
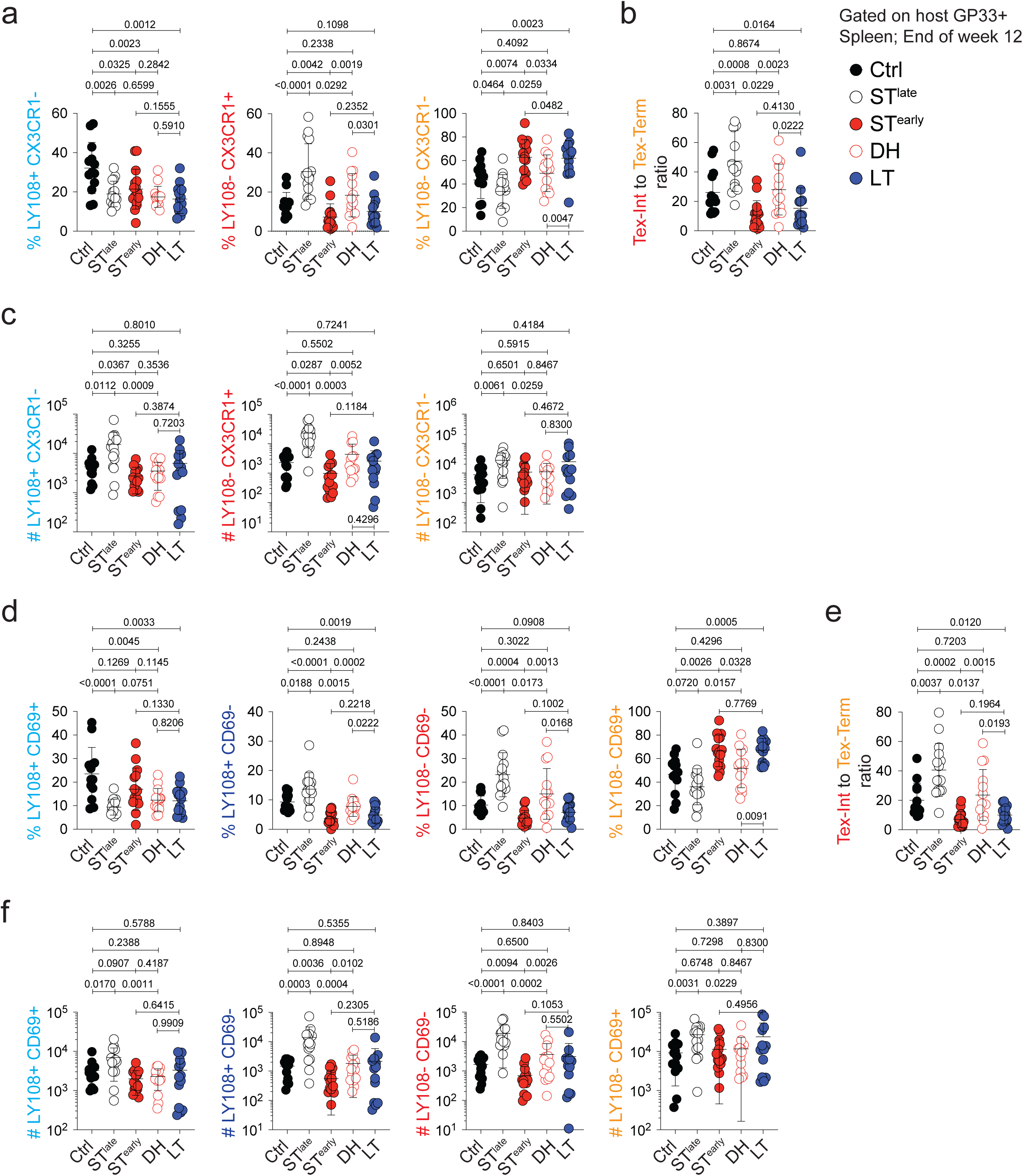
Tex subset distribution for endogenous D^b^GP_33_-tetramer+ CD8 T cells following different PD-1 blockade regimens. (a and d) Frequencies and (c and f) numbers of (a and c) LY108 and CX3CR1, and (d and f) LY108 CD69-defined Tex subsets for endogenous D^b^GP_33_-tetramer+ CD8 T cells following different PD-1 blockade regimens are shown. (b and e) Endogenous D^b^GP_33_-tetramer+ CD8 T cell (b) Tex-Int (LY108–CX3CR1+) to Tex-Term (LY108–CX3CR1–) ratios and (e) Tex-Int (LY108–CD69–) to Tex-Term (LY108–CD69+) ratios in spleen following different PD-1 blockade regimens are shown. Each dot represents an individual mouse, and error bars represent SD. Statistical significance between groups was determined by two-tailed Mann-Whitney test. (a, b, c, d, e, and f) Data shown (n = 13-15 per group) are pooled from two independent experiments.

**Extended Data Fig. 7.**
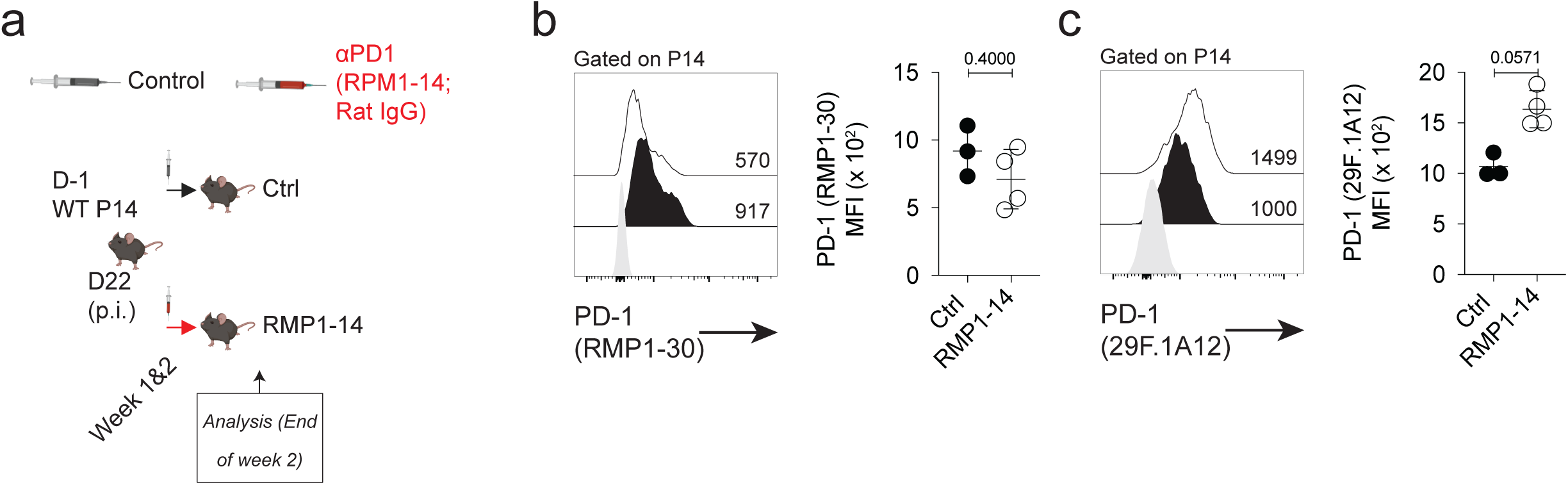
Comparison of PD-1 staining clones following anti-PD-1 (RMP1-14) blockade. Groups of LCMV clone 13-infected B6 mice containing adoptively transferred P14 cells were treated with control or anti-PD-1 (RMP1-14; rat IgG). (a) Experimental design. Representative overlaid histograms and MFI for PD-1 (b) staining clone RMP1-30 and (c) staining clone 29F.1A12 on WT P14 cells are shown. Grey histogram: gated on CD44– naïve CD8 T cells. Each dot represents an individual mouse, and error bars represent SD. Statistical significance between groups was determined by two-tailed Mann-Whitney test. Data shown (n = 3-4 per group) are from one independent experiment.

**Extended Data Fig. 8.**
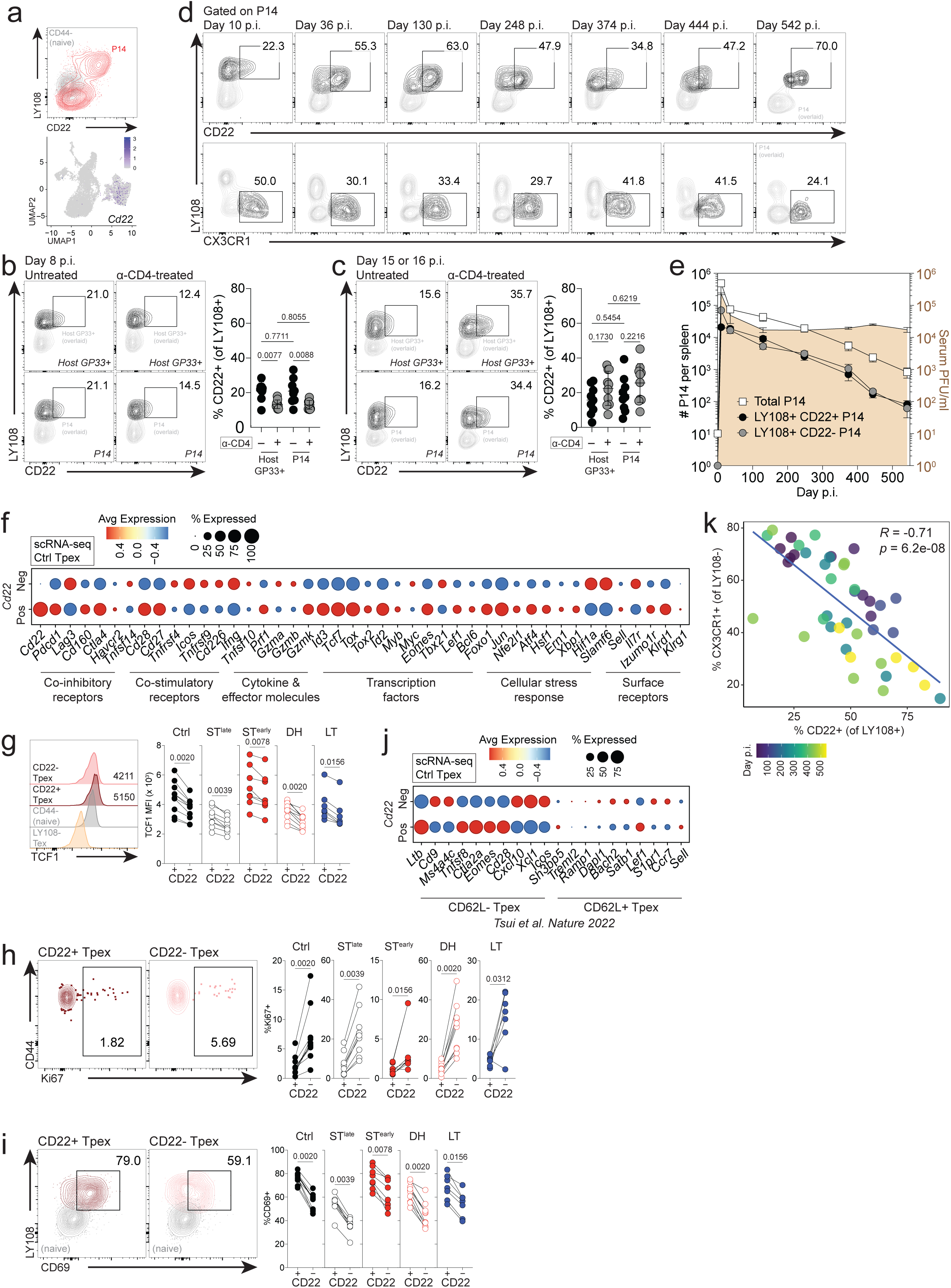
Characterization of CD22+ Tpex. (a) Representative overlaid flow cytometry plot for LY108 and CD22 expression by P14 Tex (day 111 post clone 13 infection) and CD44– naïve CD8 T cells, and *Cd22* transcript expression by P14 Tex UMAP space (as shown in Fig. 3e). (b and c) Representative overlaid flow cytometry plots for LY108 and CD22 expression for endogenous D^b^GP_33_-tetramer+ CD8 T cells and P14 Tex from (b) day 8 and (c) day 15 or 16 post LCMV clone 13 infection, in the absence or presence of anti-CD4 treatment. Summary of the proportions of CD22+ among LY108+ Tpex is shown. (d, e, and k) Mice containing P14 cells (with transient anti-CD4 treatment) were analyzed at different days post LCMV clone 13 infection. (d) Representative overlaid flow cytometry plots for LY108 and CD22 (top panel) and LY108 and CX3CR1 expression by P14 Tex are shown. (e) Summary of numerical changes for the indicated P14 cells are shown (mean ± SEM). (f and j) Ctrl-derived Tpex were defined as *Cd22*-positive and *Cd22*-negative based on scRNA-seq. (f) Dot plot illustrating the relative expression of the indicated genes in *Cd22*-positive and *Cd22*-negative Tpex is shown. (g) Representative overlaid histogram plot for TCF-1 expression in CD22+ and CD22– Tpex. (h and i) Representative flow cytometry plot and frequencies for (h) Ki67+ and (i) CD69+ between CD22+ and CD22– Tpex. (j) Dot plot illustrating the relative expression of top DEGs for CD62L+ versus CD62L- Tpex from ^36^ in *Cd22*-positive and *Cd22*-negative Tpex is shown. (k) Correlation of CD22+LY108+ cells with CX3CR1+LY108– P14 Tex is shown. (b, c, g, h, i, and k) Each dot represents an individual mouse, and (b and c) error bars represent SD. Statistical significance between groups was determined by (b and c) Kruskal-Wallis test and (g, h, and i) two-tailed Wilcoxon test. Association between CD22+LY108+ and CX3CR1+LY108– P14 Tex was calculated using (k) Pearson correlation. Data shown (b and c) (n = 9-10 per group) are pooled from two independent experiments. Data shown (d, and e) (n = 5-9 per time point), (k) (n = 44), (g, h, and i) (n = 7-10 per group) are representative of two independent experiments.

**Extended Data Fig. 9.**
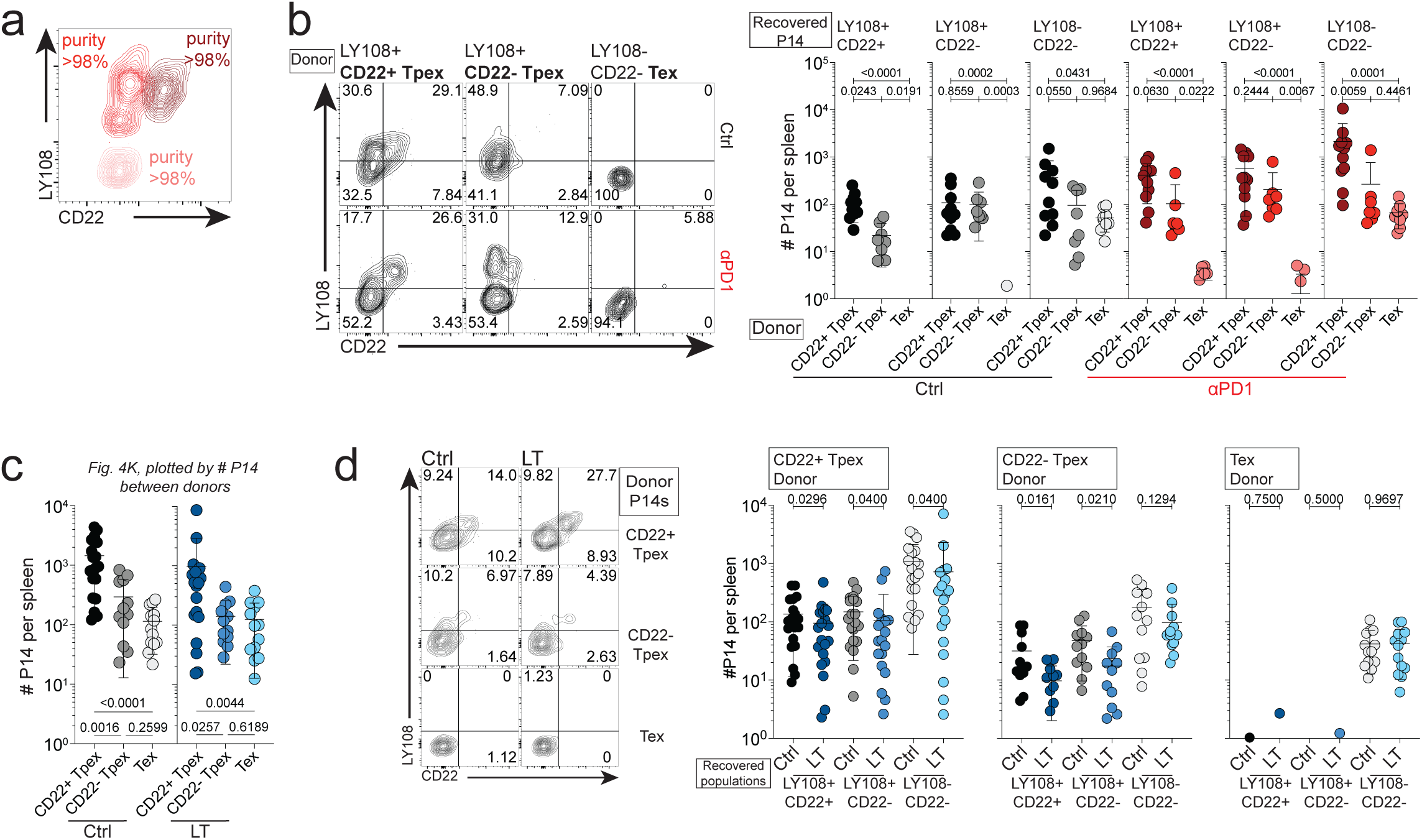
“Stemness” characterization of CD22+ Tpex. (a) Representative overlaid flow cytometry plots of sorted LY108 and CD22-defined subsets of P14 Tex for adoptive transfer experiments. (b) Representative flow cytometry plots and numbers of the indicated LY108 and CD22-defined Tex subsets in spleens of mice that received the indicated donor LY108 and CD22-defined P14 Tex subsets. (c) Fig. 4k plotted comparing Ctrl and LT donors. (d) Representative plots and numbers of the indicated LY108 and CD22-defined Tex subsets recovered in spleens of mice that received the indicated donor LY108 and CD22-defined P14 Tex subsets. Each dot represents an individual mouse, and error bars represent SD. Statistical significance between groups was determined by (b and c) Kruskal-Wallis test and (d) two-tailed Wilcoxon test. Data shown are pooled from (b) two (n = 7-11 per group) or (c and d) three (n = 12-20 per group) independent experiments.

**Extended Data Fig. 10.**
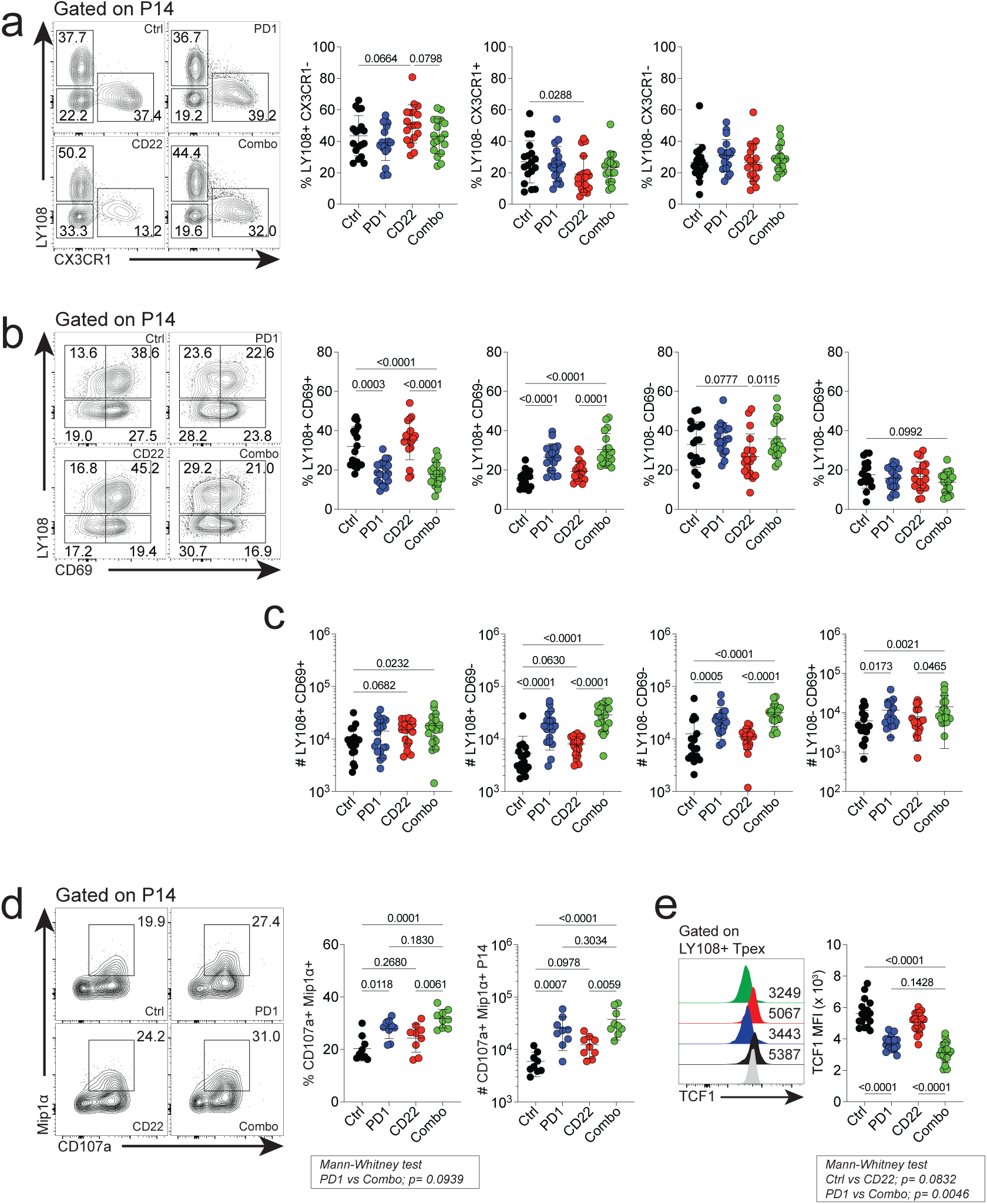
Co-blockade of CD22 and PD-1 modulates Tex subsets. (a) Representative flow cytometry plots and frequencies of LY108 and CX3CR1-defined Tex subsets in the spleen following the indicated blockade regimens are shown. (b) Representative flow cytometry plots, frequencies, and (c) numbers of LY108 and CD69-defined Tex subsets in the spleen following the indicated blockade regimens are shown. (d) Representative flow cytometry plots, frequencies, and numbers of CD107a+Mip1α+ P14 cells in the spleen following the indicated blockade regimens are shown. (e) Representative histogram plots and expression of TCF-1 in LY108+ Tpex following the indicated blockade regimens are shown. Each dot represents an individual mouse. Statistical significance between groups was determined by (a, b, c, d, and e) Kruskal-Wallis test and (d and e) two-tailed Mann-Whitney test. (a, b, c, and e) Data shown (n = 18 per group) are pooled from two independent experiments. (d) Data shown (n = 9 per group) are from one independent experiment.

**Extended Data Fig. 11.**
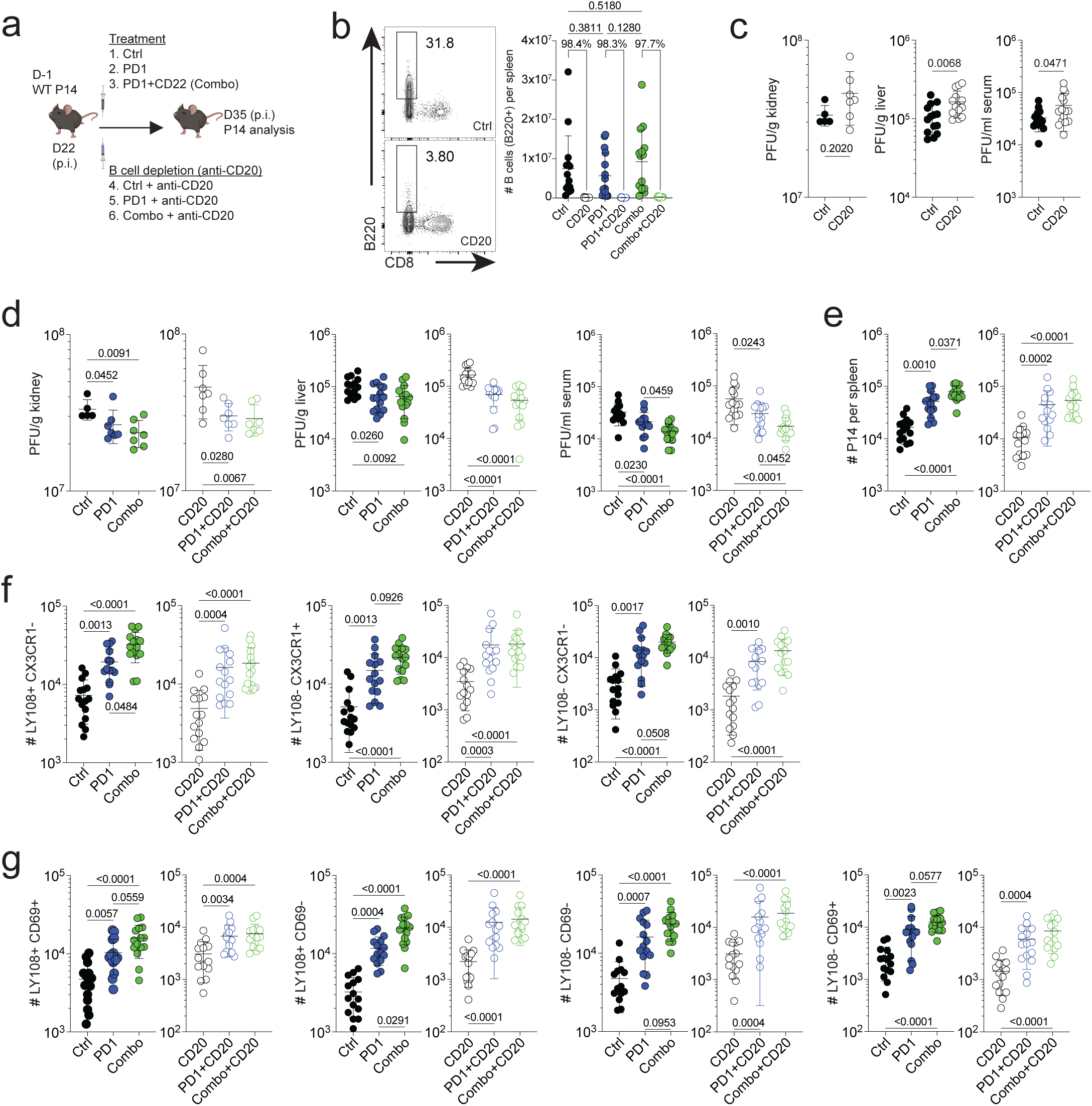
Co-blockade of CD22 and/or PD-1 in the absence of B cells. (a) Experimental design. (b) B cell depletion efficiency in anti-CD20-treated mice. (c) LCMV viral load in kidney, liver, and serum in Ctrl and anti-CD20 mice is shown. (d) LCMV viral load in kidney, liver, and serum of mice following the indicated blockade regimens is shown. (e) Numbers of P14 cells in the spleen following the indicated blockade regimens are shown. (f and g) Numbers of (f) LY108 and CX3CR1- and (g) LY108 and CD69-defined Tex subsets in the spleen following the indicated blockade regimens are shown. Each dot represents an individual mouse. Statistical significance between groups was determined by (b, d, e, f, and g) Kruskal-Wallis test and (c) two-tailed Mann-Whitney test. Data shown are (c and d; kidney panel) (n = 5-7 per group) representative of two independent experiments or (b, c, d, e, f, and g) (n = 14-15 per group) pooled from two independent experiments.

**Extended Data Fig. 12.**
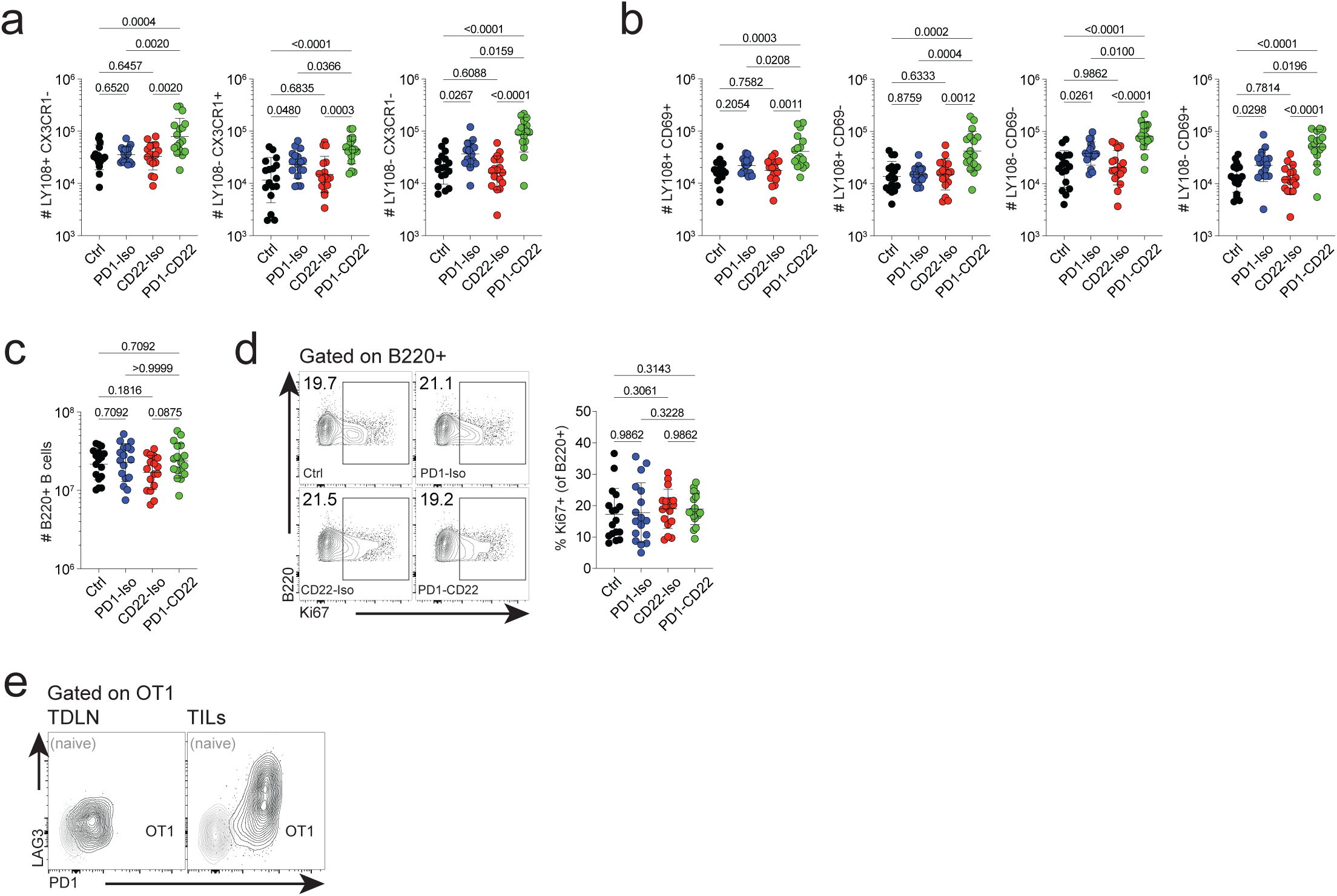
PD-1-CD22 bispecific antibody treatment modulates Tex subset biology. (a and b) Numbers of (a) LY108 and CX3CR1- and (b) LY108 and CD69-defined Tex subsets in the spleen following the indicated blockade regimens are shown. (c) Numbers of B220+ B cells in the spleen following the indicated blockade regimens are shown. (d) Representative flow cytometry plots and frequencies of Ki67+ cells as a fraction of B220+ B cells in the spleen following the indicated blockade regimens are shown. (e) Representative flow cytometry plots of PD-1 and LAG3 expression by OT-1 cells in tumor draining lymph node (TDLN) and TILs, overlaid with CD44– naive CD8 T cells (grey). (a, b, c, and d) Each dot represents an individual mouse. Statistical significance between groups was determined by (a, b, c, and d) Kruskal-Wallis test. (a, b, c, and d) Data shown (n = 17 per group) are pooled from two independent experiments.

**Supplementary Table 1.**
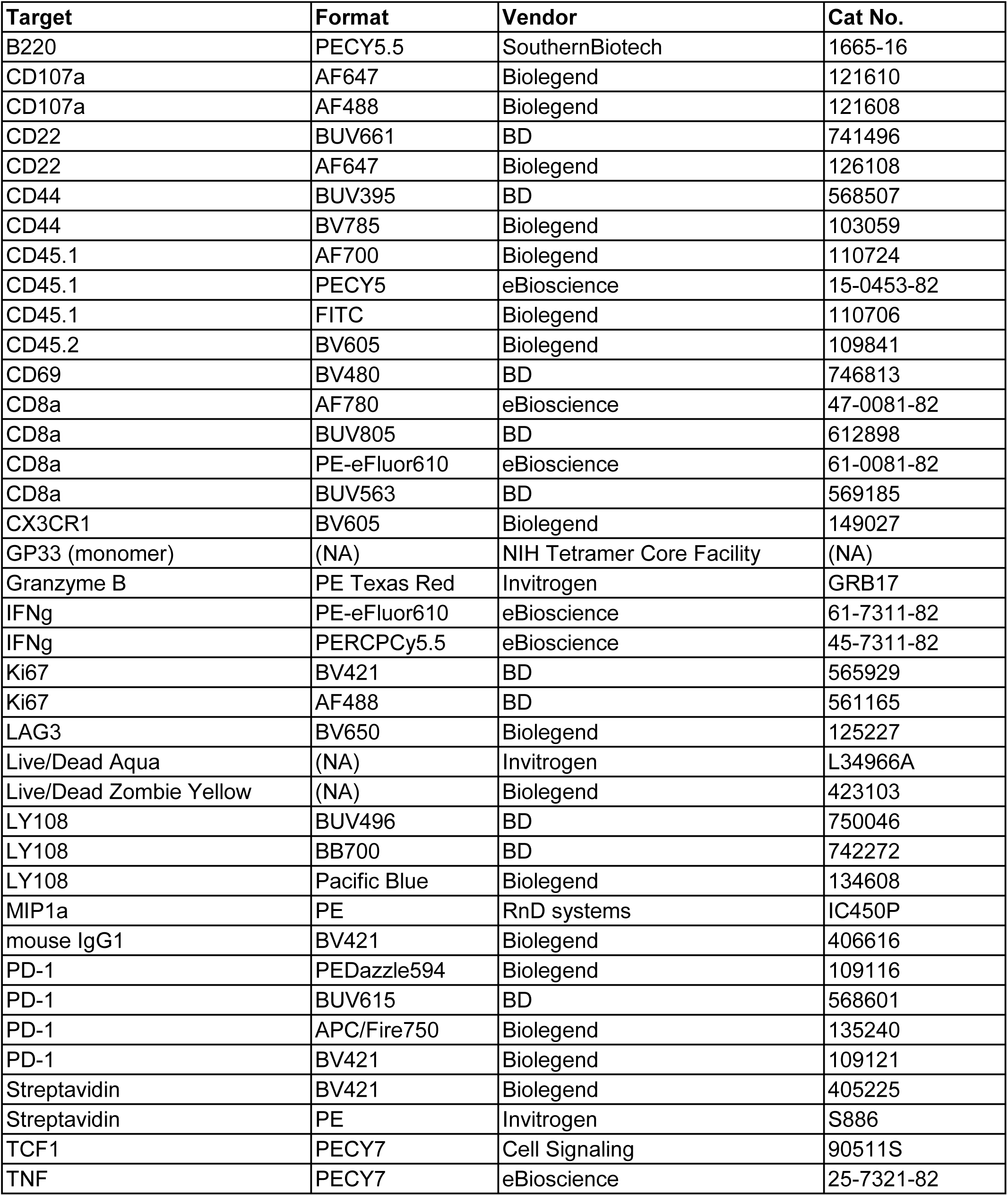
Flow cytometry targets.

